# Inflammatory bowel disease-associated gut commensals degrade components of the extracellular matrix

**DOI:** 10.1101/2022.08.09.503432

**Authors:** Ana Maria Porras, Hao Zhou, Qiaojuan Shi, Xieyue Xiao, JRI Live Cell Bank, Randy Longman, Ilana Lauren Brito

## Abstract

Extracellular matrix (ECM) remodeling has emerged as a key feature of inflammatory bowel disease (IBD), and ECM fragments have been proposed as markers of clinical disease severity. Recent studies report increased protease activity in the gut microbiota of IBD patients. Nonetheless, the relationship between gut microbiota and ECM remodeling has remained unexplored. We hypothesized that members of the human gut microbiome can degrade host ECM, and that bacteria-driven remodeling, in turn, can enhance colonic inflammation. Through a variety of *in vitro* assays, we first confirmed that multiple bacterial species found in the human gut are capable of degrading specific ECM components. Clinical stool samples obtained from ulcerative colitis patients also exhibited higher levels of proteolytic activity *in vitro* compared to those of their healthy counterparts. Furthermore, culture supernatants from bacteria species capable of degrading human ECM accelerated inflammation in a dextran sodium sulfate (DSS)-induced colitis. Finally, we identified several of the bacterial proteases and carbohydrate degrading enzymes (CAZymes) potentially responsible for ECM degradation *in vitro*. Some of these protease families and CAZymes were also found in increased abundance in a metagenomic cohort of IBD. These results demonstrate that some commensal bacteria in the gut are indeed capable of degrading components of human ECM *in vitro* and suggest this proteolytic activity may be involved in the progression of IBD. A better understanding of the relationship between nonpathogenic gut microbes, host ECM, and inflammation could be crucial to unravel some of the mechanisms underlying host-bacteria interactions in IBD and beyond.

## INTRODUCTION

Uncontrolled remodeling of the host extracellular matrix (ECM) is a known hallmark of inflammatory bowel disease (IBD) (1–6). The ECM—consisting of proteins, glycoproteins, and proteoglycans—provides not only mechanical support but also important biochemical cues for the development and homeostasis of the colon (7). Increased protease activity and degradation of the ECM in the intestinal mucosa and submucosa have been reported in both ulcerative colitis (UC) and Crohn’s disease (CD) (6, 8–11). Many IBD patients also suffer from intestinal fibrosis, which involves the accumulation of ECM components like collagen along the lining of the colonic epithelium (4, 12–14). Excessive ECM degradation and deposition may result in the development of fistulae and strictures, respectively, with serious clinical consequences (15–17). As a result, ECM fragments and proteases have emerged as potential markers of disease severity (5, 6, 18–20). Recent studies in mouse models (8, 21) and clinical settings (9, 22) suggest ECM degradation precedes inflammation in UC. Thus, dysregulated ECM production is not only a product but also a promoter of inflammation and an active player in the pathogenesis of IBD. Nonetheless, the causes of this ECM imbalance and the contributions of the gut microbiota to these dynamic ECM processes are not fully understood.

While the degradation of mucin by gut microbiota has been studied extensively (23–28), there is limited knowledge regarding the ability of commensal bacteria to degrade components of human ECM in the gut. Bacterial pathogens have been shown to bind and degrade ECM to invade intestinal and other host tissues (29–32). Similarly, bacteria associated with oral microbiota dysbiosis can break down components of the basal lamina potentially contributing to the progression of periodontal disease (33–35). Prominent members of the gut microbiome like *Bacteroides thetaiotaomicron (B. theta)* and *Bacteroides fragilis* are also known to express sulfatases (36, 37) and gelatinases (38, 39), respectively. However, the pathological consequences of this proteolytic activity have not been explored from the perspective of bacteria-ECM interactions.

We hypothesized that multiple members of the gut microbiome can remodel human ECM, and that bacteria-driven degradation, in turn, can enhance colonic inflammation. First, we designed a series of *in vitro* assays that uncovered the ability of multiple bacterial species present in the human gut to degrade various ECM components. The same assays were repeated using samples collected from healthy and UC patients. The microbiota in these clinical UC samples were more proteolytically active than those of their healthy counterparts. Finally, culture supernatants from bacteria species capable of degrading human ECM exacerbated inflammation in a mouse model of DSS-induced colitis. Collectively, the results presented in this study suggest gut microbiota indeed interact with and degrade host ECM in a manner that may contribute to the progression of IBD.

## RESULTS

### Commensal members of the gut microbiome can degrade ECM components *in vitro*

First, we performed a series of *in vitro* tests to assess the ability of commensal bacteria to degrade individual host ECM components. We selected 12 bacterial strains that are abundant in human gut microbiomes, commonly used as probiotics, and known mucin-degraders. Additionally, some of these species have previously been associated with inflammation and the progression of IBD. For example, OMVs secreted by *B. theta* are suggested to play an important role in directing immune cell behavior (36, 40) and *Ruminococcus gnavus* is commonly found in increased abundance in the microbiota of both UC and CD patients linked to disease severity (41–43). Furthermore, enterotoxigenic *B. fragilis*, found in abundance in IBD and colorectal cancer, secretes a metalloprotease capable of altering endothelial barrier integrity and inducing the secretion of inflammatory cytokines (44). These strains were cultured individually in their corresponding recommended complete growth medium (Supplementary Table 1). Because many ECM-degrading enzymes produced by pathogens are secreted (30, 45), we performed all assays using culture supernatant. Thus, supernatant from the bacterial cultures was collected and used in degradation assays for ECM components abundant in either the mucosa or submucosa (1) - collagen I and IV, laminin, fibronectin, chondroitin sulfate, and hyaluronic acid.

In these *in vitro* assays, all ECM components were degraded by components in the supernatants of at least one species (Figure 1A-F). *B. fragilis* was the primary degrader of collagen I and IV with just one other species (*Bacteroides vulgatus)* exhibiting mild proteolytic activity against these proteins (Figure 1A). In contrast, the remaining components were each degraded by supernatant from at least 3 different species (Figure 1C-F). Supernatant from a few species like *R. gnavus, B. fragilis*, and *B. theta* were particularly active in these *in vitro* degradation tests. Additionally, supernatant obtained from the genus *Bacteroides* degraded all components except for hyaluronic acid. In contrast, we detected little to no proteolytic activity in species often proposed as probiotics (46, 47) like *Lactobacillus gasseri, Lactobacillus reuteri*, and *Bifidobacterium longum* (Fig 1A-F).

**Figure 1.**
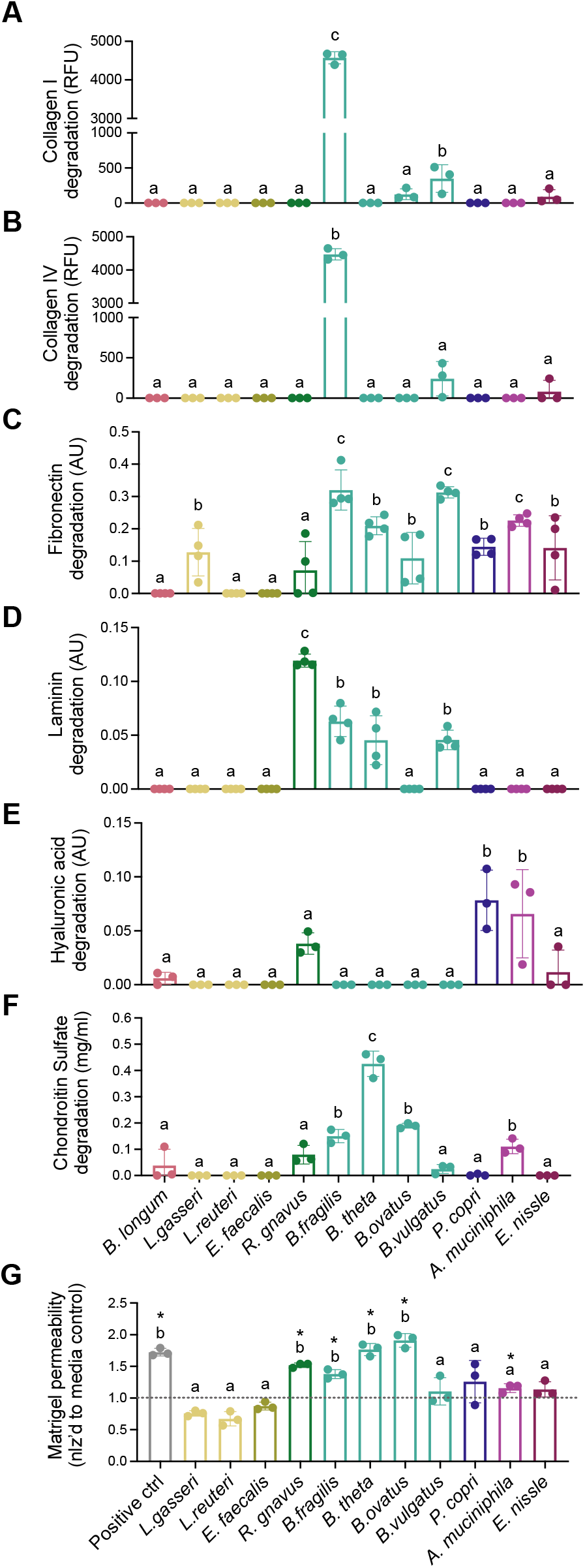
Commensal members of the gut microbiome can degrade ECM components *in vitro*. (A-F) *In* vitro degradation of (A) collagen I, (B) collagen IV, (C) fibronectin, (D) laminin, (E) hyaluronic acid, and (F) chondroitin sulfate by supernatant obtained from the individual culture of 12 bacterial species present in the human gut microbiome. Species represented with the same color belong to the same phylum. (G) Permeability of a Matrigel-based *in vitro* model of the basement membrane after 24 hours of culture with bacterial supernatant. For all panels, n = 3-4 replicated and data are presented as mean ± SD. Same letters denote groups that are not statistically different; different letters indicate groups that are statistically different from each other, *p* < 0.05 by one-way ANOVA followed by Tukey’s multiple comparison test.

We then developed a Matrigel-based model of the basement membrane to test ECM degradation using a more complex substrate. Bacterial culture supernatant supplemented with FITC-labeled dextran was added to the top of a trans-well insert pre-coated with a Matrigel layer. Matrigel permeability after 24 hours of incubation was then assessed by measuring fluorescence at the bottom of the well. As observed in the other *in vitro* assays, incubation with supernatant from *R. gnavus* and bacteria from the genus *Bacteroides* genus (*B. fragilis, B. theta, and B. ovatus)* led to significantly higher permeability compared to media-only controls (Figure 1G, one-way ANOVA with Tukey’s multiple comparison test). This was not surprising considering most of these species had previously exhibited proteolytic activity against collagen IV and laminin, two of the most abundant components of Matrigel and the basement membrane.

To complement our findings, we also evaluated strain- and isolate-level differences using the same *in vitro* assays (Figure 2). We selected two additional clinical specimens of *B. fragilis* strains (ATCC 43858 and DSM 9669) for comparison against the type strain (ATCC 25285, Supplementary Table 1). Additionally, we included three *Prevotella copri* isolates obtained from a participant in the FijiCOMP project (48) for comparison against the type strain (DSM 18205). For most of the ECM components evaluated, there were statistically significant differences between additional strains and isolates and the corresponding type strain (Figure 2; one-way ANOVA with Tukey’s multiple comparison test). For example, the *B. fragilis* type strain degraded collagen I, collagen IV, and chondroitin sulfate to a greater extent than either of the other strains (Figures 2A-B, 2F). In the case of laminin (Figure 2D) and hyaluronic acid (Figure 2E), the *P. copri* and *B. fragilis* type strains, respectively, exhibited no enzymatic activity while the isolates and commensal strains were indeed capable of breaking down these components. These results highlight the importance of considering strain-level differences in microbiome studies.

**Figure 2.**
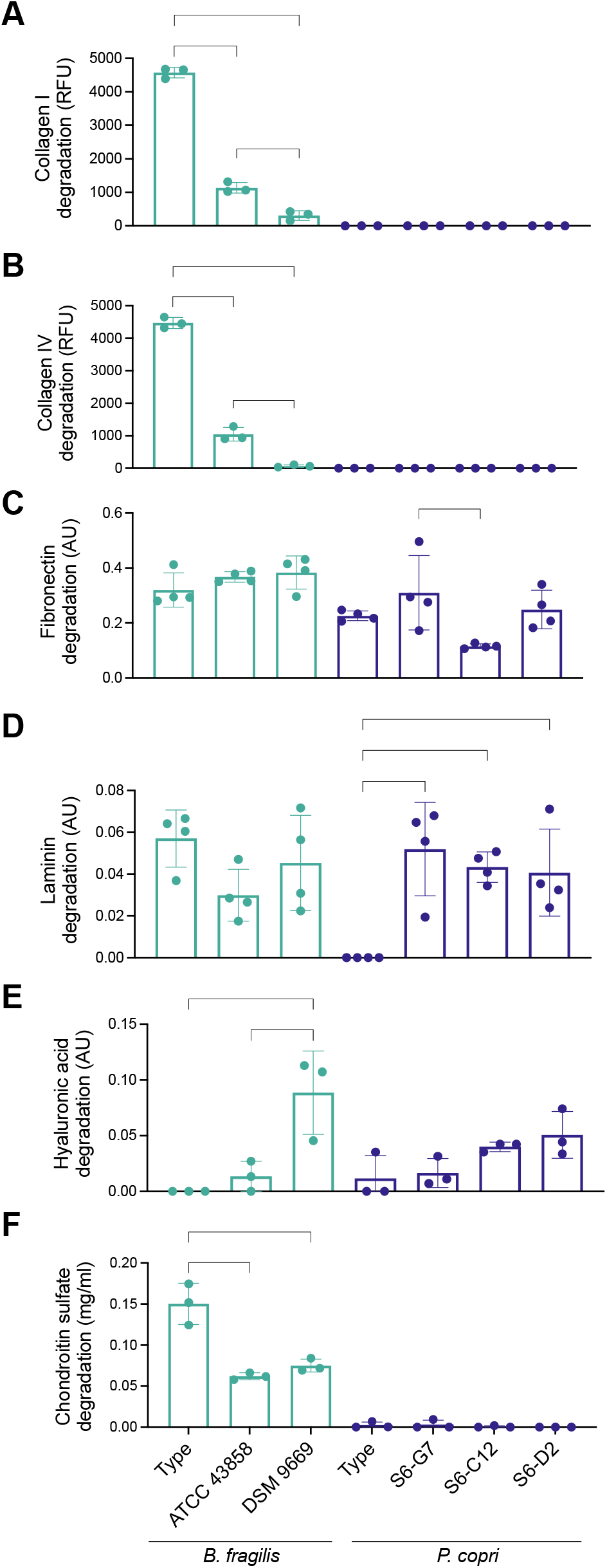
Strains and isolates of the same bacterial species exhibit differences in ECM degradation *in vitro*. (A-F) *In* vitro degradation of (A) collagen I, (B) collagen IV, (C) fibronectin, (D) laminin, (E) hyaluronic acid, and (F) chondroitin sulfate by supernatant from *B. fragilis* strains and *P. copri* isolates. Bars of the same color indicate the same species. For all panels, n = 3-4 replicated and data are presented as mean ± SD. *p<0.05, **p<0.01, ***p<0.001, ****p<0.0001 by one-way ANOVA followed by Tukey’s multiple comparison test.

### Supernatant from clinical ulcerative colitis samples exhibits higher proteolytic activity

Next, we assessed the capacity of stool community supernatants to degrade ECM components in a clinical context using the *in vitro* assays described above. We included 19 samples from healthy (n=10) and UC (n=9) patients (Supplementary Table 2). These samples were stored at -80°C and resuspended in prereduced PBS supplemented with cysteine in an anaerobic chamber to create a stock solution. This stock solution was then inoculated at 2% (v/v) in two culture media – supplemented Brain Heart Infusion Broth (BHIS) or Gut Microbiome Medium (GMM) (49). Because the use of any one medium would lead to the preferential growth of some microorganisms, we instead opted to use two different culture media. BHIS was selected given its ability to support the growth of *Bacteroides* species – the most proteolytically active in the *in vitro* assays (Figure 1). GMM, on the other hand, was employed because of its reported ability to support the growth of a wide diversity of bacteria compared to other media (49). Supernatant from these cultures was collected 24 hours after inoculation and subjected to the same *in vitro* single-component ECM degradation assays described before.

In general, the supernatant obtained from UC patients were better able to degrade individual ECM substrates compared to their healthy counterparts (Figure 3; two-way ANOVA with Tukey’s multiple comparison test). More specifically, the UC samples exhibited increased proteolytic activity against (Figure 3A), collagen IV (Figure 3B), fibronectin (Figure 3C), and laminin (Figure 3D). No statistically significant differences in chondroitin sulfate degradation were observed (Figure 3E). Similarly, no statistically significant differences in the degradation of individual ECM components were observed when comparing the BHIS and GMM growth conditions for each patient group (Figure 3A-E). We also evaluated Matrigel permeability in the basement membrane model following incubation with patient supernatant for 24 hours. As expected from previous results, incubation with UC supernatant led to higher permeability compared to healthy supernatant (Figure 3F). In this case, we did observe statistically significant differences between the BHIS and GMM growth conditions in the UC group.

**Figure 3.**
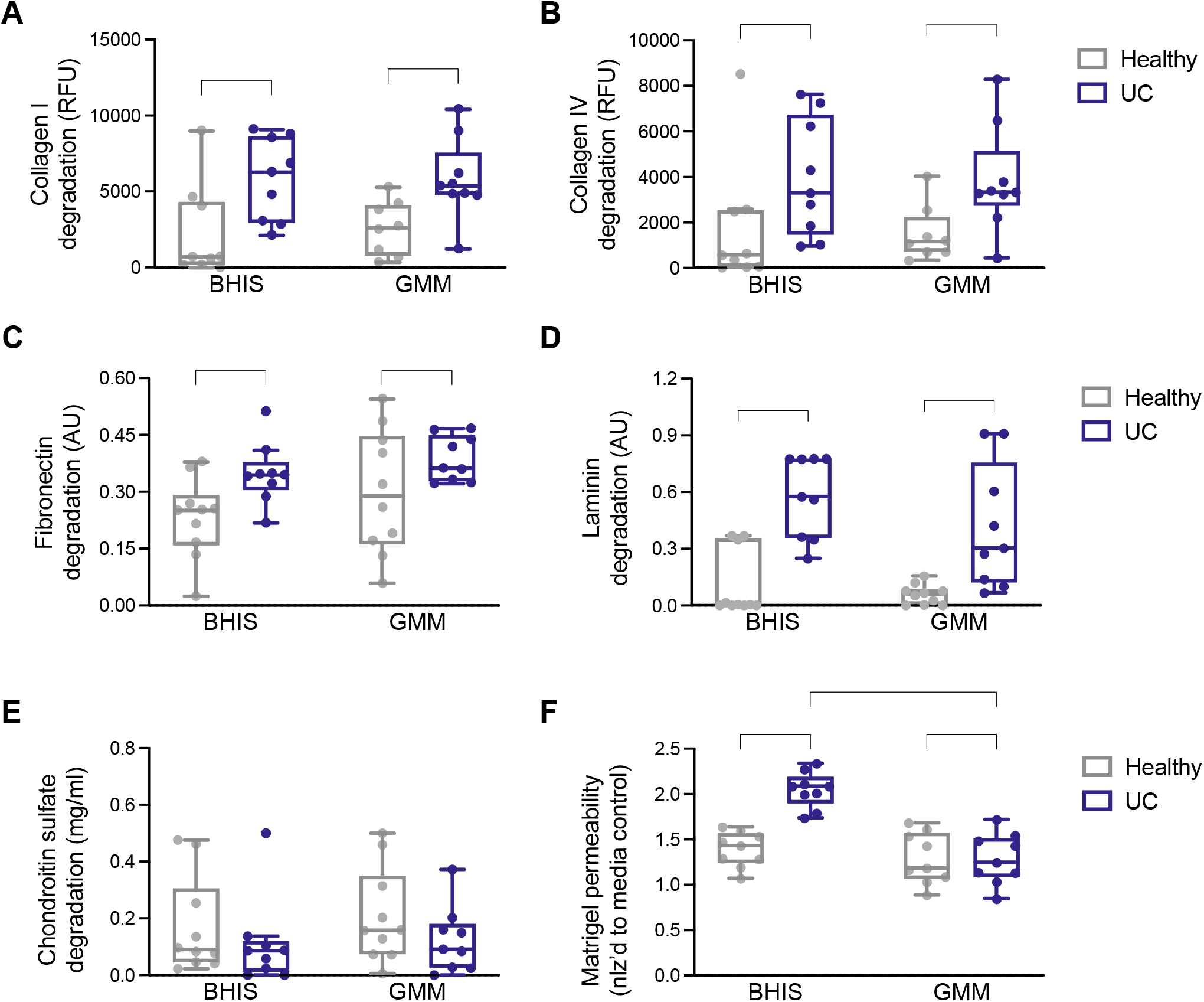
Supernatant from clinical ulcerative colitis samples exhibits higher proteolytic activity. Samples obtained from UC and healthy patients were cultured for 24 hours in BHIS or GMM. Culture supernatant was then subjected to a variety of ECM degradation assays. (A-E) *In* vitro degradation of (A) collagen I, (B) collagen IV, (C) fibronectin, (D) laminin, and (E) chondroitin sulfate by supernatant from UC and healthy patient microbiota cultures. (F) Permeability of a Matrigel-based *in vitro* model of the basement membrane after 24 hours of culture with supernatant from UC and healthy patient microbiota cultures. For all panels, n = 9-10 and data are presented as mean ± SD. *p<0.05, **p<0.01, ***p<0.001, ****p<0.0001 by two-way ANOVA followed by Tukey’s multiple comparison test.

We performed 16S rRNA sequencing on the patient microbiomes cultured in BHIS and GMM and used to test their degradative qualities (Supplementary Figure 1). Although there were compositional differences between microbiomes grown in BHIS and GMM, the average Bray-Curtis difference was smaller between individuals’ samples grown in the two conditions, versus between individuals grown in the same medium (Supplementary Figure 1), suggesting that any bias due to media choice preserved the identity of the sample. Despite the ability for cultured microbiomes to degrade ECM components and Matrigel, we were only able to detect 3 species in the cultured microbiomes: *B. fragilis* (5 healthy; 5 UC), *A. muciniphila* (1 healthy; 1 UC) and *R. gnavus* (1 UC). Despite the degradative qualities of *B. fragilis*, it’s abundances after culture were higher overall in the healthy samples (Supplementary Figure 1). This highlights the likelihood that the degradative traits are common across a broader subset of species.

### Exposure to proteolytic supernatants accelerates inflammation in a DSS-induced mouse model of IBD

We also explored the effects of repeated exposure to bacterial supernatants in a dextran sulfate sodium salt (DSS)-induced colitis mouse model. Specifically, we selected supernatant from three of the most proteolytically active species in the *in vitro* assays – *B. fragilis* (ATCC 43858), *B. theta* and *R. gnavus*. C57BL/6 mice were treated with 1.5% DSS in drinking water for 10 consecutive days to induce acute colitis. Mice were gavaged daily with either bacterial supernatant or culture medium before, during, and after DSS (Figure 4A, n = 9 mice per treatment group). Weight loss in all DSS-treated groups, regardless of exposure to supernatants, became significant by day 8 post-DSS treatment peaking around 15% weight loss by day 10 with no statistically significant differences between groups (Figure S2).

**Figure 4.**
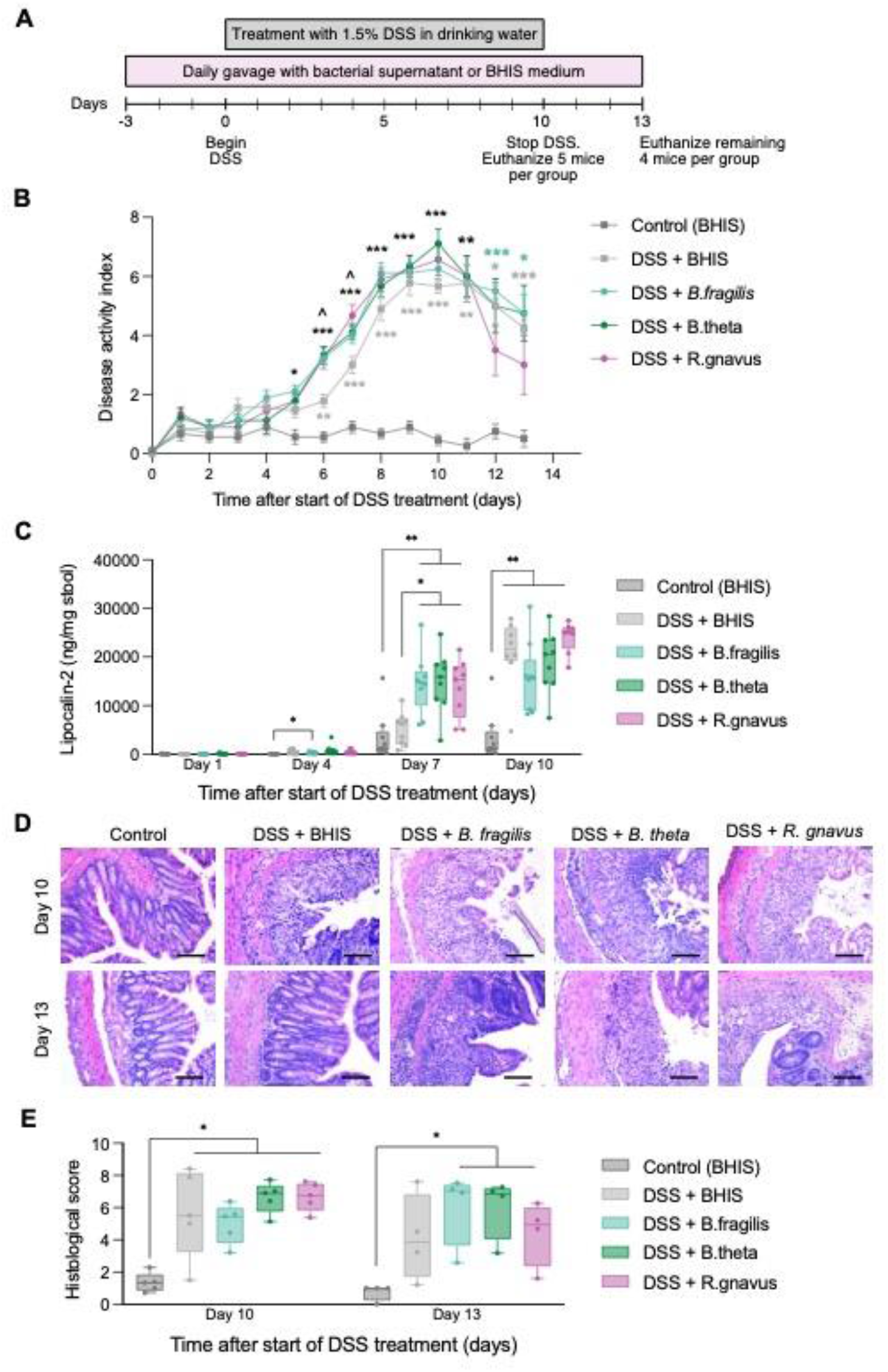
Exposure to proteolytic supernatants accelerates inflammation in a DSS-induced mouse model of IBD. (A) Schematic of the *in vivo* experimental set up. (B) Disease activity index over time after the start of DSS treatment. Data represent mean ± SD. *p<0.05, **p<0.01, and ***p<0.001 compared to Control (BHIS). ^p<0.05 compared to DSS + BHIS. Asterisks in black indicate all supernatant experimental groups achieved that level of significance. (C) Quantification of lipocalin-2 levels in mouse stool at days 1,4, 7, and 10 post-DSS treatment. *p<0.05 and **p<0.01 for comparisons shown. (D) H&E-stained cross sections of explanted mouse colons on days 10 and 13. Scale bar represents 100 μm. (E) Histological score quantifying the colonic tissue damage observed in (D). *p<0.05 for comparisons shown. For all panels, statistical significance was assessed using a mixed-effects model with the Geisser-Greenhouse correction followed by Tukey’s multiple comparisons test.

There were, however, differences in the timing of onset of symptoms. Analysis of the disease activity index (DAI) curve showed equally severe clinical symptoms between DSS-only (also treated with BHIS media as a control) and supernatant-treated groups by day 8 post DSS-treatment (Figure 4B); however, all supernatant-treated mice began exhibiting clinical signs of colitis earlier than the DSS-only group (at day 5 rather than day 6). Additionally, the DAI also increased more rapidly in the supernatant-treated mice between days 5 and 8. The amount of lipocalin-2 (LCN-2) in the stool, a clinical biomarker of inflammatory diseases (50), was significantly elevated in all DSS + supernatant-treated mice by day 7 compared to the control and DSS-only groups (Figure 4C; mixed-effects model with the Geisser-Greenhouse correction followed by Tukey’s multiple comparisons test). By day 10, LCN-2 levels were elevated in all groups that received DSS treatment.

Mice treated with supernatant were also slower to recover than DSS-only treated mice. We noticed that by day 10, all DSS-treated groups exhibited extensive epithelial damage, crypt ablation, and mucosal erosion as observed by histology, compared to the untreated control group (Figures 4D-4E). Greater variability in the extent of tissue damage was observed in the DSS-only group compared to all other experimental groups. However, on day 13, 3 days after ending DSS treatment, the colons of the DSS –only mice showed some evidence of recovery and improved tissue architecture (Figure 4D). By contrast, the mice that continue to receive daily gavage with *B. fragilis, B. theta*, and *R. gnavus* supernatant sustained statistically significant tissue damage, crypt destruction, and immune cell infiltration (Figures 4D-E; mixed-effects model with the Geisser-Greenhouse correction followed by Tukey’s multiple comparisons test). Overall, these results suggest that daily gavage with proteolytically active supernatant may accelerate inflammation and sustain tissue damage *in vivo*.

### General proteases are identified in bacterial culture supernatants and overexpressed in IBD clinical cohorts

Finally, we sought to identify the proteases and carbohydrate degrading enzymes (CAZymes) found in each strain’s supernatant that could be responsible for degradation of the ECM components analyzed. We performed untargeted proteomic analysis of culture supernatant obtained from the species that exhibited significant proteolytic behavior against any of the assayed ECM components. These strains included *A. muciniphila, B. fragilis* (type, ATCC 43858, and DSM 9669), *B. ovatus, B. theta, B. vulgatus, P. copri* (Type, S6-G7, S6-C12, and S6-D2), and *R. gnavus*. After annotating the protein families (Pfams) and CAZymes secreted by each strain, we identified those previously reported to play a role in the degradation of ECM generally or of specific components (*e*.*g*., hyaluronic acid or laminin). These curated protein families included multiple metalloproteases (M18 and M12B), as well, as several general proteases known to degrade a variety of ECM components like trypsin, papain, and calpain (Supplementary Table 3). We also identified multiple CAZymes involved in the degradation of proteoglycans and glycosaminoglycans like alpha-amylase, beta-xylosidase, and both alpha- and beta-mannosidases (Supplementary Table 4).

To confirm the clinical relevance of these enzymes, we assessed their relative abundances in the PRISM dataset, a cohort that includes healthy, UC, and Crohn’s disease patients (51). Thirteen out of the 47 enzymes on the curated list were differentially-abundant between healthy and IBD patients (Figure 5; Mann-Whitney U-test with false discovery rate correction). Among Pfams, trypsin, peptidase families M23 and U32, and PPIC-type PPIase domain were found at a higher relative abundance in IBD microbiomes compared to the healthy controls (Figure 5A-E). Similarly, N-acetylglucosamine deacetylase, β-galactosidase, β-glucosidase, chitinase, and α-mannosidase (Figure 5G-L) were also more abundant in the IBD samples. In contrast, both the M18 zinc metalloprotease (Figure 5F) and β-xylosidase (Figure 5M) were relatively less abundant in IBD versus healthy microbiomes. These results implicate bacteria-secreted proteases and CAZymes in the process of degrading ECM leading to the progression of IBD.

**Figure 5.**
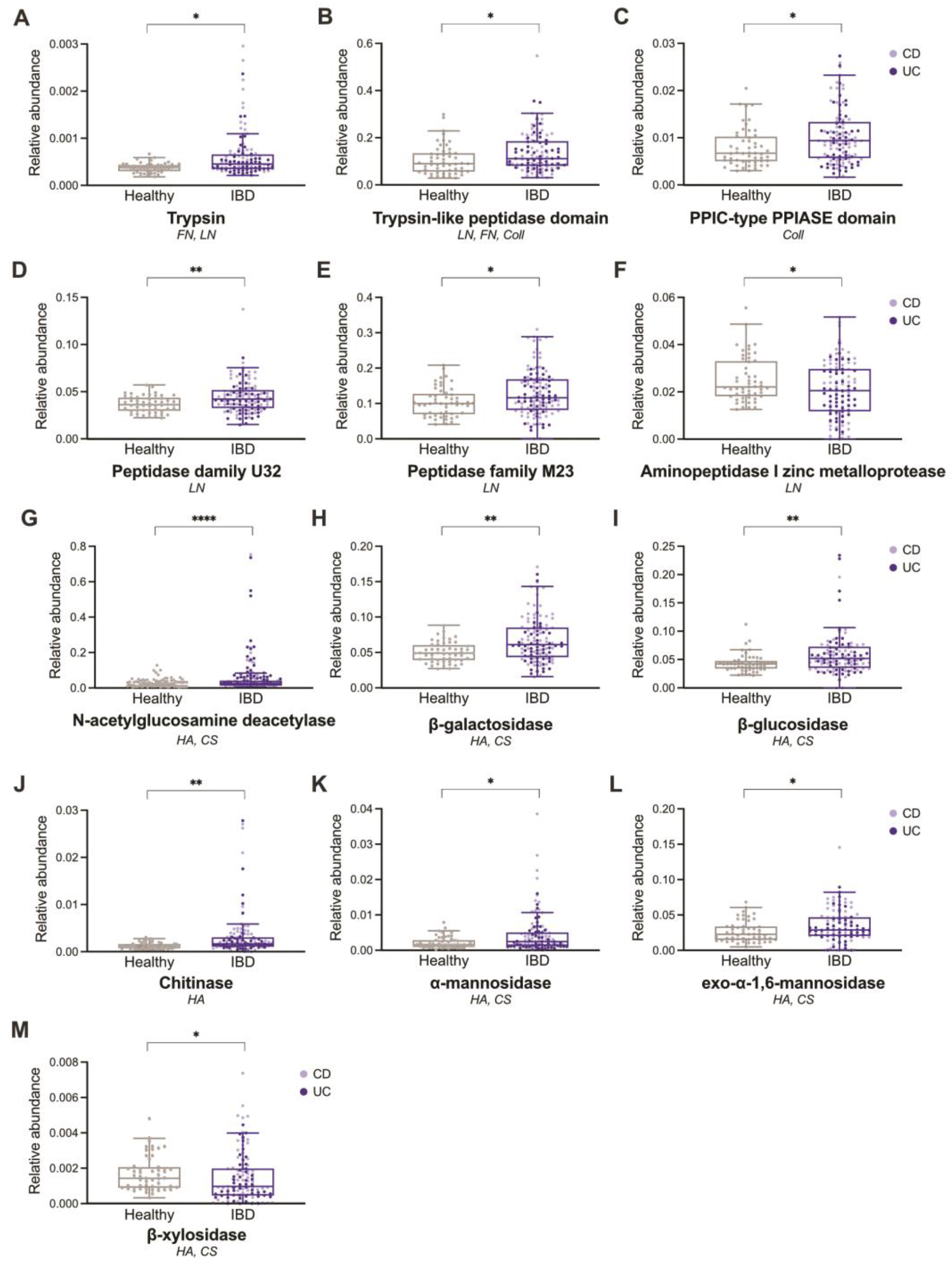
Proteases and CAZymes secreted by ECM-degrading bacterial strains *in vitro* are differentially abundant in an IBD cohort compared to healthy controls. Relative abundance of the protein families (A-F) and CAZymes (G-M) found to be significantly different between IBD and healthy metagenomes from the PRISM dataset (51). For each protein and enzyme family, the substrates degraded by the strain supernatans in which they were detected *in vitro* are listed. FN = fibronectin, Coll – collagen I and IV, LN = laminin, HA = hyaluronic acid, CS = chondroitin sulfate. For all panels, statistical significance was calculated using Mann-Whitney U-test with false discovery rate correction, where *p<0.05, **p<0.01, ***p<0.001.

## DISCUSSION

ECM remodeling is increasingly recognized as a key step in the progression of disease and a potential therapeutic target for IBD (9, 52, 53). Mounting evidence points to increased activity of fecal (19, 22, 54) and, specifically, bacterial proteases (55, 56) associated with disease severity in UC. While these studies link bacterial proteolytic activity to inflammation, the specific mechanisms involved have not been identified. Here, we demonstrate that commensal gut microbiota secrete proteases and CAZymes capable of ECM degradation *in vitro*. Several commensal bacteria were particularly good ECM degraders including several species of the genus *Bacteroides, R. gnavus*, and *P. copri*. In some cases, specific strains of *R. gnavus and B. fragilis* contribute unevenly to IBD pathophysiology (57, 58). We extend these observations, showing differences between the ECM degradation capabilities of strains of *B. fragilis* and *P. copri*.

We specifically identified serine and cysteine proteases, metalloproteinases, and glycosyl hydrolases, in the bacterial supernatants exhibiting the highest proteolytic activity *in vitro*. Of these, we found trypsin, and several metalloproteases (peptidase families U32 and M23, and aminopeptidase I zinc) in increased abundance in a large metagenomic IBD cohort. Elevated serine and trypsin-like protease activity has also been reported in other analyses of IBD fecal samples (54, 55, 59) with increasing evidence that these enzymes are secreted by commensal bacteria of the *Bacteroides* genus (22, 55, 60). Similarly, zinc-dependent metalloproteases secreted by pathogenic bacteria contribute to the deterioration of intestinal barrier function through a variety of mechanisms primarily targeting endothelial cells (61). Our results point to a role for these proteases secreted by gut commensals in the degradation of multiple ECM proteins including collagen, laminin, and fibronectin. Moreover, they also highlight the importance of CAZymes like α-mannosidase, β-galactosidase, and β-glucosidase not only in the digestion of food and mucin (62), but also in the breakdown of glycosaminoglycans and glycoproteins commonly found in gut ECM.

Secretion of these enzymes by commensals is unlikely to induce IBD on its own. In a healthy gut, the gut microbiota is confined to the intestinal lumen by a thick layer of mucus and would therefore not have access to the underlying ECM (63). In contrast, in IBD a variety of genetic and environmental factors can disrupt the balance between mucosal barrier and gut microbiota. Steck *et al*. demonstrated that the matrix metalloprotease gelatinase E, secreted by *E. faecalis*, can degrade E-cadherin and induce inflammation in a disease susceptible *IL-10*^*-/-*^ mouse background but not in wild-type mice (64). Furthermore, disruption of the endothelial membrane in UC and CD patients leads to invasion of colonic tissue by bacteria, including those of the genus *Bacteroides* (65, 66). Our data suggests that alterations of intestinal homeostasis could provide an opportunity for commensal-derived proteases to encounter host ECM and induce tissue damage.

ECM degradation by commensal microbiota can lead to serious consequences for the host. In this study, exposure to proteolytic supernatant in a DSS-induced model of colitis accelerated the manifestation of inflammation symptoms and led to an increase in Lipocalin-2 levels. Shimshoni *et al*. recently demonstrated ECM degradation precedes symptoms of inflammation in a similar mouse model (21). There are a variety of mechanisms through which bacteria-driven ECM remodeling can contribute to the progression of IBD. First, degradation of components of the basement membrane like collagen IV and laminin could further disrupt epithelial integrity (6, 67). Second, degradation of submucosal ECM can precipitate the recruitment and activation of immune cells (52, 68). For example, cleavage of hyaluronic acid (69, 70) and collagen (8) triggers the recruitment of nearby leukocytes.

Our results identify a potential role for gut microbiota in host ECM remodeling, and, as a result, IBD progression. Microbiome-sourced proteases and CAZymes may serve as potential drug targets to ameliorate damage to the ECM in IBD, although additional work is necessary to determine the potential success of such a treatment. It will be necessary to determine the relative contributions of host- and microbial-derived metalloproteases and to determine the specificity of each bacterial enzyme in order to predict their effects on the host. Additionally, larger cohorts will be necessary in order to establish whether the abundance or total activity of these enzymes correlates with disease severity, and most importantly, ECM-associated phenotypes, *i*.*e*. fibrotic lesions. Finally, it remains to be determined how secreted bacterial enzymes gain access to the extracellular matrix, preceding overt damage to the epithelial cell lining. Nevertheless, our work provides additional mechanistic understanding of the roles that IBD-associated bacteria play in this disease.

## METHODS

### Bacterial Culture

Bacterial strains were grown anaerobically at 37°C in an anaerobic chamber (COY Lab Products) in their corresponding complete growth medium as outlined in Supplementary Table 1. *P. copri* strains were isolated from samples collected as part of the Fiji Community Microbiome Project (FijiCOMP) (47). This study was initially approved by the Institutional Review Boards at Columbia University, the Massachusetts Institute of Technology, and the Broad Institute and ethics approvals were received from the Research Ethics Review Committees at the Fiji National University and the Ministry of Health in the Fiji Islands. The Cornell University Institute Review Board additionally approved this study (#1608006528). Human subjects were consented prior to participation in the study. To prepare supernatants, liquid cultures were inoculated from frozen glycerol stocks and grown to an OD_600_ of 1.0 to 1.1. At that point, cultures were centrifuged at 7,000 x g for 10 minutes and the supernatant was collected and separated from the bacterial pellets. The supernatant was then refrigerated at 4°C for a maximum of 6 hours until all supernatants were ready to begin the ECM degradation assays.

### Quantification of ECM Degradation *in vitro*

Specific degradation tests were selected for each ECM component. In all cases, background degradation levels were considered based on the corresponding culture medium for each bacterial supernatant. A SpectraMax (Molecular Devices) plate reader was used to measure fluorescence and absorbance for all assays.

Gelatin and collagen degradation were quantified using the EnzChek Gelatinase/Collagenase Assay Kit (Thermo Fisher). DQ gelatin, collagen I, or collagen IV were added to bacterial supernatants and media controls at a final concentration of 50 μg/mL. Fluorescence (abs. 495 nm/ em. 515 nm) was measured in the solution following overnight incubation at 37°C under anaerobic conditions. In this case, fluorescence was directly proportional to gelatin and collagen degradation.

Fibronectin and laminin degradation were instead evaluated using a modified ELISA protocol adapted from work by Mendes *et al*. (71). High binding 96 well-plates were coated with recombinant fibronectin (2 μg/mL; Millipore Sigma) and laminin (1 μg/mL; Millipore Sigma) diluted in PBS and incubated overnight at 37°C. The next day, plates were washed 3 times with 1x PBS and blocked with 3% BSA in PBS-T for at least 2 hours at 37°C. After removing the BSA, bacterial supernatant was added to the corresponding wells in quadruplicates and the plate. After a second anaerobic overnight incubation at 37°C, the plates were washed 3 times with 1x PBS to remove the supernatant and the degradation of the pre-coated ECM components was detected using mouse anti-fibronectin (F7387, 1:5,000; Millipore Sigma), and rabbit anti-laminin (L9393, 1:10,000; Millipore Sigma) antibodies diluted in PBS-T for 1 hour at 37°C. Following another series of washes, HRP-conjugated goat anti-rabbit IgG (1:5,000; Millipore Sigma) and anti-mouse IgG (L9393, 1:5,000; Millipore Sigma) were added to the plates for 1 hour at 37°C. Finally, TMB-ELISA substrate solution (Thermo Fisher) was added and the reaction was stopped with 2N H_2_SO_4_. Absorbance in this case was inversely proportional to protein degradation.

HA degradation was analyzed in a similar fashion (72). High binding 96 well-plates were coated with 200 μg/mL HA (Millipore Sigma) diluted in 0.2 M carbonate buffer, pH 9.2 and incubated overnight at 4°C. Following washing with 1x PBS, non-specific binding was blocked 3% BSA in PBS-T for at least 2 hours at 37°C. The plates were washed and supernatant was added to the corresponding wells and incubated anaerobically overnight at 37°C. The HA remaining after supernatant-driven degradation was detected with HA binding protein following dilutions and instructions in the Hyaluronan DuoSet ELISA kit by R&D Systems (DY3614). As was the case for fibronectin and laminin, absorbance levels were inversely proportional to protein degradation.

The degradation of chondroitin sulfate (CS) was evaluated using a quantitative Alcian Blue assay. A 10 mg/ml stock solution of CS from shark cartilage (Millipore Sigma) was prepared in deionized water. That stock was then diluted to a final concentration of 0.5 mg/mL in bacterial supernatant and incubated anaerobically at 37°C overnight. Alcian Blue dye stock was prepared by diluting 0.5 g of Alcian Blue (VWR) in 100 mL of 18 mM H_2_SO_4_, centrifuging the solution at 10,000 x g for 30 mins, and filtering. CS standards ranging from 2 to 0.004 mg/mL were prepared in deionized water. After the overnight incubation, 10 μL of standards or sample were added to a microcentrifuge tube followed by 10μL of sample diluent (4 M guanidine containing 0.0375% Triton X-100 in 27 mM H_2_SO_4_) and 100 μL of working Alcian Blue solution (5% dye stock in 18 mM H_2_SO_4_ + 0.25% Triton X-100). The tubes containing samples or standards were then vortexed briefly to mix, and centrifuged at 10,000 x g for 10 min. The supernatant was then decanted and the pellets were left to dry for at least 10min. Finally, the pellets were dissolved in 100 μL of 8 M guanidine by vortexing. The solutions were then pipetted into a 96-well plate and absorbance was read at 600 nm. The generation of a standard curve allowed us to quantify the final CS concentration in each sample after supernatant-driven degradation.

### Matrigel-Based Basement Membrane Degradation Model

We designed an additional degradation model that better captured the complexity of the basement membrane based on a tissue penetration model described by Andrian *et al*. (73). Matrigel (Corning) was diluted 1:3 in cold PBS and 100 μL was added to 0.4 μm polycarbonate trans-well plate inserts (VWR). The Transwell plates were placed 4°C for 30 min to let the Matrigel settle and were then moved to an anaerobic chamber to gel at 37°C for 24 h. The next day, Matrigel was rehydrated in 100 μL of sterile reduced PBS for 1 h at 37°C. In the meantime, a 10 mg/mL stock of 40 kDa FITC-labeled Dextran (Millipore Sigma) was prepared and later diluted in either media or supernatant at a final concentration of 0.5 mg/mL. 150 μL of the supernatant containing FITC-labeled Dextran was pipetted on top of the Matrigel and 300 μL of PBS were added to the lower chamber. The Transwell plates were then incubated anaerobically for 24h at 37°C. Fluorescence in the bottom chamber was measured to assess permeability of the Matrigel layer. Because dextran is a carbohydrate that could be digested by gut bacteria, the percent of dextran that successfully traversed the membrane was calculated in comparison to the fluorescence levels in leftover FITC-dextran and supernatant solution after the same anaerobic incubation for 24 h at 37°C.

### Preparation of Human Stool Supernatants

Stool microbiome samples were obtained from informed and consented patients during colonic irrigation procedures in accordance with IRB protocols for Weill Cornell Medical College (#1501015812) and Cornell University (#1609006586). Ulcerative colitis was defined by clinical or endoscopic characteristics. Healthy samples were collected on 2017-2019. Between 0.5-1mL of sample were frozen after collection and moved to storage at -80°C. To prepare stool stocks for culture, stool was resuspended in pre-reduced PBS supplemented with 0.05% L-cysteine-HCL to make a stock solution. Frozen stool stocks were inoculated at a concentration of 2% (v/v) in 5mL of either BHIS or Gut Microbiome Medium (GMM) (49). Liquid cultures were grown overnight for 24 hours and the supernatant was collected after centrifugation at 7,000 x g for 10 minutes. Immediately after, the proteolytic activity of these stool culture supernatants was assessed through the ECM degradation assays described above.

### DSS-Induced Colitis Mouse Model

*B. fragilis, B. theta* and *R. gnavus* supernatants were prepared by growing up 25 ml of culture overnight in BHIS medium, centrifugation at 7,000 rpm for 10 mins, and filtering through a 0.22 μm Steriflip (EMD Millipore) filter unit. The supernatants were then frozen at -80°C in 2 mL aliquots. Aliquots of BHIS medium were also prepared.

This mouse study was performed following protocols approved by the Cornell Institutional Animal Care and Use Committee (Protocol ID #2016-0088). 45 male SPF C57BL/6 mice (The Jackson Laboratory) at 7 weeks of age were obtained for this experiment and housed individually during treatments. After one week of acclimatization, we started treating the mice with the supernatants through daily oral gavage (200 μL per mice). Nine mice were treated per bacterial strain with two additional control groups receiving daily gavages of BHIS medium. On the 4^th^ day of treatment with supernatant, acute ulcerative colitis was induced by exposure to 1.5% (w/v) dextran sulfate sodium salt (DSS, 36,000-50,000 M.Wt., MP Bio) in drinking water *ad libitum* for 10 consecutive days for all mice except for one of the BHIS groups that received normal drinking water. Fresh water with DSS was replaced every 3 days. Five mice per group were sacrificed at the end of DSS treatment. Following the end of DSS treatment, the daily gavage with supernatant continued for another 3 days until the remaining mice were sacrificed. Fecal pellets were collected daily. Mice were monitored for weight loss, food and water intake, pathological features (rectal bleeding and diarrhea), and survival. They were also inspected for visible clinical signs of pathology. The presence of diarrhea, rectal bleeding, and weight lost were separately graded on a 0 to 3 scale (Supplementary Table 5) adapted from Gommeaux *et al*. (74). The scores were then added to calculate the disease activity index (DAI).

### Histological and Immunofluorescent Characterization of Explanted Mouse Colons

Sections (0.2-0.5 cm) of the terminal colon were collected after euthanasia, fixed in either formalin or methacarn for 24 hours, and later placed in 70% or 100% ethanol, respectively. The tissue sections were then paraffin embedded and sectioned at the Animal Health Diagnostic Center at the Cornell University College of Veterinary Medicine, where H&E staining was also performed on formalin-fixed sections. Antigen retrieval was performed prior to immunofluorescent staining by heating the tissue sections in citric acid buffer (pH 6.0; Vector Laboratories) at 95°C for 20 minutes. The sections were then washed with PBS and blocked with 10% goat serum overnight followed by another overnight incubation at 4°C with monoclonal antibodies against laminin (1:200; Sigma Aldrich, L9393) and collagen IV (1:400, Abcam, ab6586) diluted in 1% goat serum in PBS. After washing with PBS, secondary goat anti-rabbit IgG Alexa Fluor antibodies (Thermo Fisher, A1108) were applied diluted 1:500 in 1% goat serum in PBS. Finally, coverslips were mounted with ProLong Gold Antifade Mountant with DAPI (Thermo Fisher). Colorimetric and fluorescent images were obtained on an inverted Leica DMi8 microscope. H&E Images were blinded prior to histopathological scoring and we used the method described by Bonfiglio *et al*. (75) to quantitatively describe lamina propria cellularity, architectural damage, and epithelial abnormalities (Supplementary Table 5).

### Quantification of Lipocalin-2 in Mouse Stool

We followed the protocol by Chassaing *et al* to asses lipocalin-2 levels in the mouse stool (50). Stool pellets were reconstituted in PBS containing 0.1% Tween 20 (100 mg stool/mL) followed by 10 mins of vortexing and centrifugation at 10,000 x g for 10 mins. The supernatant was then collected and frozen at -20°C. LCN-2 levels were quantified later by ELISA (DY1851, R&D Systems).

### 16s rRNA Gene Sequencing

We extracted genomic DNA from human stool cultures or mouse fecal pellets using QIAGEN DNeasy PowerSoil kits following the manufacturer’s instructions. The V4 region of the 16S rRNA gene was amplified in triplicate following Earth Microbiome Project protocols (76), and using barcoded 515F (77) and 806R (78) primers, and the Platinum Hot Start PCR Master Mix (Thermo Fisher). PCR products were cleaned using AMPure XP beads and pooled for each sample. Prior to sequencing, amplicon pools were quantified with Quant-iT PicoGreen dsDNA Assay Kit (Invitrogen). 100 ng of amplicons from each sample were pooled prior to submission for paired-end sequencing on the Illumina MiSeq platform at the Cornell Institute of Biotechnology.

16S rRNA gene sequences were analyzed using the Quantitative Insights into Microbial Ecology (QIIME2; https://qiime2.org/) pipeline. First, we performed quality control with DADA2 (80) to remove chimeric sequences, retain unique sequence variants, and trim forward and reverse reads. Taxonomies were assigned using QIIME2’s Naïve Bayes classifier trained with the Greengenes Database). We then used the scipy.spatial.distance.braycurtis function to compute Bray-Curtis Distance.

### Supernatant preparation for nano LC/MS/MS

Following bacterial culture as described above, the supernatant was collected and filtered using a 10 KDa Amicon ultra-4 centrifugal filter unit (Millipore Sigma) at 15,000 x g and 4°C for 15 minutes. The supernatant was then concentrated 10-fold in PBS containing SIGMA*FAST* protease inhibitor (Millipore Sigma), frozen at -20°C and submitted to the Proteomics Facility at the Cornell Institute of Biotechnology. In solution digestion for each sample was performed with a S-Trap micro spin column (ProtiFi, Huntington, NY, USA) following a Strap protocol as described previously (79, 80) with slight modifications. 30 micrograms of proteins in 25 µL buffer containing 50 mM TEAB pH 8.5, 6 M Urea, 2 M Thiourea, 1% SDS were reduced with 15 mM Dithiothreitol (DTT) for 1 h at 34 °C, alkylated with 50 mM iodoacetamide for 1 h in dark and then quenched with a final concentration of 25 mM DTT. After quenching, 12% phosphoric acid was added to each sample for a final concentration of 1.2%, followed by 1:7 dilution (v/v) with 90% methanol, 0.1 M TEAB pH 8.5. Each of the resulting samples was then placed into a spin column and centrifuged 3000 x g for 30 sec. Then washed three times with 150 µl 90% methanol, 0.1 M TEAB pH 8.5. Digestion was performed by adding 25 µl trypsin at 1:10 w/w (trypsin: proteins) in 50 mM TEAB pH 8.5 to the top of the spin column. The spin columns were incubated overnight (16 hr) at 37 °C. Following incubation, the digested peptides were eluted off the S-trap column sequentially with 40 µl each of 50 mM TEAB pH 8.5 followed by 0.2% formic acid and finally, 50% acetonitrile, 0.2% formic acid. Three eluates with eluted peptides were pooled together and evaporated to dryness by a Speedvac SC110 (Thermo Savant, Milford, MA).

### Identification of Proteins in Bacterial Supernatants

The tryptic digests were reconstituted in 0.5% formic acid (FA) for nanoLC-ESI-MS/MS analysis. The analysis was carried out using an Orbitrap Fusion™ Tribrid™ (Thermo-Fisher Scientific, San Jose, CA) mass spectrometer equipped with a nanospray Flex Ion Source, and coupled with a Dionex UltiMate 3000 RSLCnano system (Thermo, Sunnyvale, CA) (79, 81). The peptide samples (20μL) were injected onto a PepMap C-18 RP nano trapping column (5 µm, 100 µm i.d x 20 mm) at 20 µL/min flow rate for rapid sample loading and then separated on a PepMap C-18 RP nano column (2 µm, 75 µm x 25 cm) at 35 °C. The tryptic peptides were eluted in a 60 min gradient of 7% to 38% ACN in 0.1% formic acid at 300 nL/min., followed by a 7 min ramping to 90% ACN-0.1% FA and an 8 min hold at 90% ACN-0.1% FA. The column was re-equilibrated with 0.1% FA for 25 min prior to the next run. The Orbitrap Fusion was operated in positive ion mode with spray voltage set at 1.9 kV and source temperature at 275°C. External calibration for FT, IT and quadrupole mass analyzers was performed. Data-dependent acquisition (DDA) mode was used for analysis. The instrument was operated using FT mass analyzer during MS scan to select precursor ions followed by 3 second “Top Speed” data-dependent CID ion trap MS/MS scans at 1.6 m/z quadrupole isolation for precursor peptides with multiple charged ions above a threshold ion count of 10,000 and normalized collision energy of 30%. MS survey scans set at a resolving power of 120,000 (fwhm at m/z 200), for the mass range of m/z 375-1575. Dynamic exclusion parameters were set at 50 s of exclusion duration with ±10 ppm exclusion mass width. All data were acquired using Xcalibur 4.4 operation software (Thermo Fisher Scientific).

Peptides were identified against the corresponding genomes downloaded from the NCBI RefSeq Database (Supplementary Table 1). Open reading frames were predicted using Prodigal v2.6.3 (82). The resulting coding sequences were annotated by aligning to the Carbohydrate Active Enzyme database (http://www.cazy.org/; (83)) using DIAMOND blastp (identity >= 40%; coverage >80%; e-value < 1e-5) (84). Protein families were annotated on the Pfam-A 33.1 database using Hmmsearch v3.1 (85). For every ECM component, we compiled a list of the Pfams and CAZymes secreted by the species capable of degrading that component. We then manually inspected all Pfams and CAZymes to identify those reported to be associated with or capable of ECM degradation (Supplementary Tables 3-4).

### Analysis of Proteases and CAZymes in IBD Cohorts

We downloaded the PRISM dataset (51) and removed samples with abnormally low (less than 10^7) reads. Low-quality reads were removed using Trimmomatic-0.3 (86). We used HUMAnN3 to define the functional potential of the gut metagenome with default settings. As described in the previous section, we generated a list of protein families and CAZymes secreted by the bacterial species in the *in vitro* experiments associated with ECM degradation. We searched this curated list against Uniref90 groups identified in the PRISM dataset using DIAMOND blastp, requiring greater than 50% sequence identity and greater than 80% coverage. For each sample, we aggregated the abundances of Uniref90 groups according to corresponding protein families. Fold change differences were compared by Mann-Whitney U-test with false discovery rate (FDR) correction (FDR<0.05).

### Statistical Analysis

Statistical analysis for all experiments was performed using GraphPad Prism v9 except for the analysis of protease and CAZyme abundance in the IBD metagenomic cohort. For all strain-level experiments, groups were compared using one-way ANOVA followed by Tukey’s multiple comparison test. Two-way ANOVA followed by Tukey’s multiple comparison test was selected for the evaluation of human clinical samples. Finally, the *in vivo* data was analyzed using a mixed effects that took into account repeated measures over time with the Geisser-Greenhouse correction followed by Tukey’s multiple comparison test. In all cases, two experimental groups were considered to be statistically significant when the P value was less than 0.05 after multiple comparison corrections.

## ACKNOWLEDGEMENTS

We thank the Proteomics and Metabolomics Facility of Cornell University for providing the mass spectrometry data and NIH SIG grant 1S10 OD017992-01 support for the Orbitrap Fusion mass spectrometer. This work was supported by funding from the Cornell Presidential Postdoctoral Fellowship (to A.M.P.) and a Seed Grant from Cornell’s Institute of Biotechnology (to I.L.B.). I.L.B. is also supported by a Packard Foundation Fellowship, an NIH New Innovator Award (1DP2HL141007) and a Pew Foundation Fellowship.

## AUTHOR CONTRIBUTIONS

Conceptualization, A.M.P. and I.L.B; Resources, R.L. and JRI Live Cell Bank; Methodology, A.M.P. and I.L.B.; Investigation, A.M.P., Q.S., and X.X.; Data Curation, Formal Analysis, and Visualization, A.M.P. and H.Z.; Project administration, A.M.P. and I.L.B.; Supervision, I.L.B.; Writing – original draft, A.M.P., H.Z., and I.L.B.; Writing – review and editing, A.M.P., H.Z., Q.S., X.X., R.L., and I.L.B.

## SUPPLEMENTARY DATA

**Supplementary Table 1.** List of bacterial strains tested in degradation assays *in vitro*.

| Bacterial species | Strain | Source | Growth Medium |
| --- | --- | --- | --- |
| *Akkermansia muciniphila* | Muc | DSM 22959 | PYG Medium with 0.5% mucin |
| *Bacteroides fragilis* | NCTC 9343 | ATCC 25285 | Supplemented Brain Heart Infusion Broth |
| *Bacteroides fragilis* | 2-078382-3 | ATCC 43858 | Supplemented Brain Heart Infusion Broth |
| *Bacteroides fragilis* | MPRL 1842 | DSM 9669 | Supplemented Brain Heart Infusion Broth |
| *Bacteroides thetaiotaomicron* | VPI 5482 | DSM 2079 | Supplemented Brain Heart Infusion Broth |
| *Bacteroides ovatus* | NCTC 11153 | ATCC 8483 | Supplemented Brain Heart Infusion Broth |
| *Bacteroides vulgatus* | NCTC 11154 | ATCC 8482 | Supplemented Brain Heart Infusion Broth |
| *Bifidobacterium longum* | S12 | ATCC 15697 | ATCC Medium 2107: Modified Reinforced Clostridial |
| *Enterococcus faecalis* | Tissier | DSM 20478 | Trypticase Soy Yeast Extract Medium |
| *Escherichia coli* | Nissle 1917 |  | Luria Broth |
| *Lactobacillus gasseri* | F 164 | DSM 20077 | MRS Medium |
| *Lactobacillus reuteri* | MM4-1A | ATCC PTA-6475 | MRS Medium |
| *Ruminococcus gnavus* | H2_28 | DSM 108212 | PYG Medium |
| *Prevotella copri* | CB7 | DSM 18205 | BBL™ Schaedler Broth |
| *Prevotella copri* | S6-G7 isolate | Isolated from Fijian donor | BBL™ Schaedler Broth |
| *Prevotella copri* | S6-C12 isolate | Isolated from Fijian donor | BBL™ Schaedler Broth |
| *Prevotella copri* | S6-D12 isolate | Isolated from Fijian donor | BBL™ Schaedler Broth |

**Supplementary Table 2.**
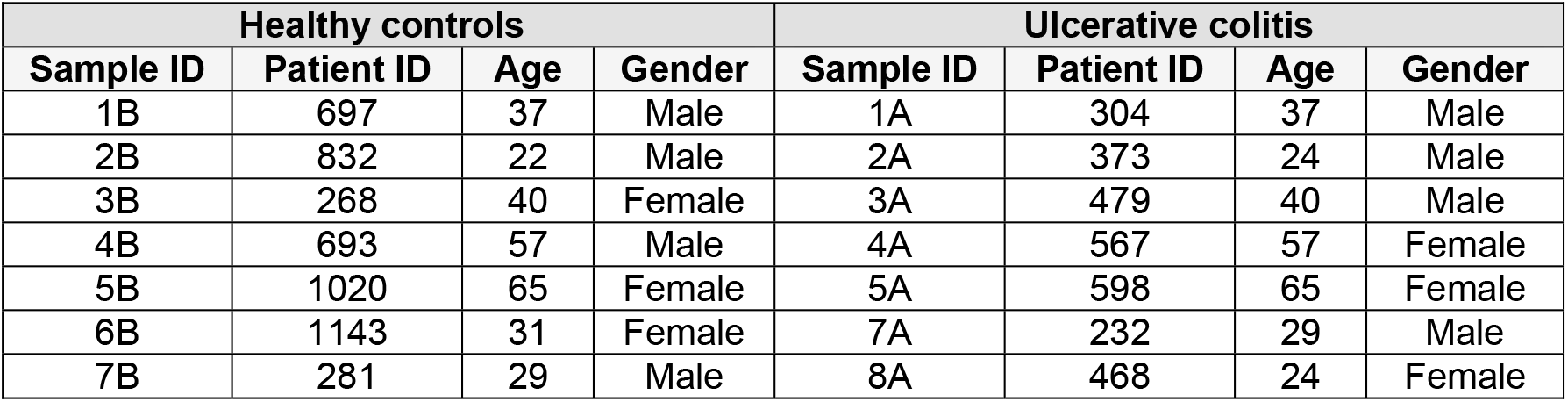
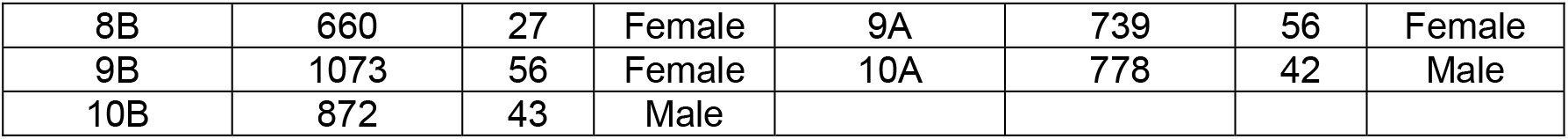
Metadata of healthy and UC patients participating in this study.

| Healthy controls |  |  |  | Ulcerative colitis |  |  |  |
| --- | --- | --- | --- | --- | --- | --- | --- |
| Sample ID | Patient ID | Age | Gender | Sample ID | Patient ID | Age | Gender |
| 1B | 697 | 37 | Male | 1A | 304 | 37 | Male |
| 2B | 832 | 22 | Male | 2A | 373 | 24 | Male |
| 3B | 268 | 40 | Female | 3A | 479 | 40 | Male |
| 4B | 693 | 57 | Male | 4A | 567 | 57 | Female |
| 5B | 1020 | 65 | Female | 5A | 598 | 65 | Female |
| 6B | 1143 | 31 | Female | 7A | 232 | 29 | Male |
| 7B | 281 | 29 | Male | 8A | 468 | 24 | Female |
| 8B | 660 | 27 | Female | 9A | 739 | 56 | Female |
| 9B | 1073 | 56 | Female | 10A | 778 | 42 | Male |
| 10B | 872 | 43 | Male |  |  |  |  |

**Supplementary Table 3.** Protein families (Pfams) associated with ECM degradation secreted by bacterial strains *in vitro*. List of Pfams secreted by bacterial species *in vitro* with reported roles involved in the degradation of ECM components. The “ECM Component” column indicates that Pfam was identified in the supernatant of bacteria capable of degrading that particular component.

| Pfam ID | Name | ECM Component |
| --- | --- | --- |
| PF01120.19 | Alpha-L-fucosidase | Laminin, collagen |
| PF02127.17 | Aminopeptidase I zinc metalloprotease (M18) | Laminin |
| PF01400.26 | Astacin (Peptidase family M12A) | Laminin |
| PF00648.23 | Calpain family cysteine protease | Fibronectin, laminin |
| PF13620.8 | Carboxypeptidase regulatory-like domain | Laminin |
| PF08669.13 | Glycine cleavage T-protein C-terminal barrel domain | Laminin, collagen |
| PF03065.17 | Glycosyl hydrolase family 57 | Fibronectin |
| PF14509.8 | Glycosyl-hydrolase 97 C-terminal, oligomerisation | Laminin |
| PF17829.3 | Gylcosyl hydrolase family 115 C-terminal domain | Laminin |
| PF02275.20 | Linear amide C-N hydrolases, choloylglycine hydrolase family | Laminin |
| PF13582.8 | Metallo-peptidase family M12B Reprolysin-like | Fibronectin, laminin, collagen |
| PF00112.25 | Papain family cysteine protease | Fibronectin, laminin |
| PF01640.19 | Peptidase C10 family | Fibronectin, laminin |
| PF01364.20 | Peptidase family C25 | Fibronectin, laminin |
| PF03577.17 | Peptidase family C69 | Fibronectin |
| PF01551.24 | Peptidase family M23 | Laminin |
| PF03571.17 | Peptidase family M49 | Laminin |
| PF01136.21 | Peptidase family U32 | Laminin |
| PF00639.23 | PPIC-type PPIASE domain | Collagen |
| PF16141.7 | Putative glycoside hydrolase Family 18, chitinase_18 | Laminin |
| PF00082.24 | Subtilase family | Laminin |
| PF00089.28 | Trypsin | Fibronectin, laminin |
| PF13365.8 | Trypsin-like peptidase domain | Laminin, collagen, fibronectin |

**Supplementary Table 4.** CAZymes associated with glycosaminoglycan degradation secreted by bacterial strains *in vitro*. List of CAZymes secreted by bacterial species *in vitro* with reported roles involved in the degradation of glycosaminoglycans (in this case, HA and CS).

| CAZy ID | Functions |
| --- | --- |
| CE5 | acetyl xylan esterase |
| GH57 | alpha-amylase, alpha-galactosidase |
| GH13 | alpha-amylase, pullulanase |
| GH31 | alpha-glucosidase, alpha-galactosidase, alpha-mannosidase, alpha-xylosidase |
| GH29 | alpha-L-fucosidase; alpha-1,3/1,4-L-fucosidase |
| GH38 | alpha-mannosidase |
| GH109 | alpha-N-acetylgalactosaminidase |
| GH2 | beta-galactosidase, beta-mannosidase |
| GH35 | beta-galactosidase, exo-beta-glucosaminidase |
| GH3 | beta-glucosidase, xylan 1,4-beta-xylosidase, beta-glucosylceramidase |
| GH20 | beta-hexosaminidase, lacto-N-biosidase, beta-1,6-N-acetylglucosaminidase |
| GH120 | beta-xylosidase |
| GH43 | beta-xylosidase, alpha-L-arabinofuranosidase, xylanase |
| GH18 | chitinase, lysozyme |
| GH101 | endo-alpha-N-acetylgalactosaminidase |
| GH51 | endoglucanase, endo-beta-1,4-xylanase, beta-xylosidase |
| GH125 | exo-alpha-1,6-mannosidase |
| GH97 | glucoamylase, alpha-glucosidase, alpha-galactosidase |
| GH0 | glycoside hydrolases not yet assigned to a family. |
| GH92 | mannosyl-oligosaccharide alpha-1,2-mannosidase |
| GH84 | N-acetyl beta-glucosaminidase, hyaluronidase |
| CE9 | N-acetylglucosamine 6-phosphate deacetylase |

**Supplementary Table 5.** Criteria for scoring the disease activity index (DAI).

| Score | Weight lost (% of initial) | Stool consistency | Rectal bleeding |
| --- | --- | --- | --- |
| 0 | <1 | Normally formed pellets | None |
| 1 | 1-4.99 | Soft pellets not adhering to the anus | Small spots of blood in stool; dry anal region |
| 2 | 5-10 | Very soft pellets adhering to the anus | Large spots of blood in stool; blood appears through anal orifice |
| 3 | >10 | Liquid stool on long streams; wet anus | Deep red stool; blood spreads largely around the anus |

**Supplementary Figure 1.**
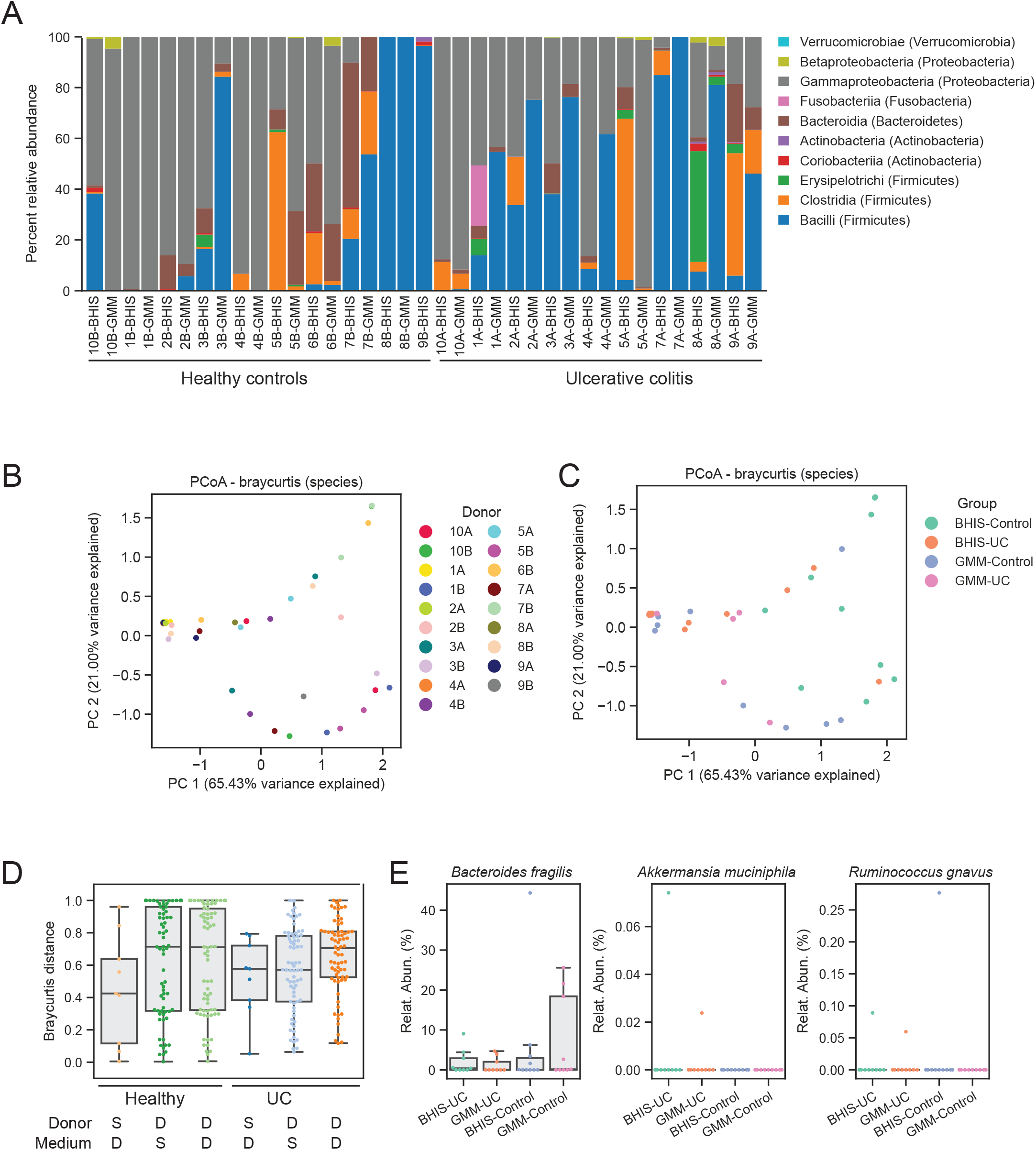
Individuals’ microbiome compositions retain similarities across media. (A) Relative abundance calculations for individuals’ microbiome samples after culture in BHIS medium or GMM. (B) A Principal Coordinate Analysis (PCoA) of samples’ species abundances. Samples are color coded according to the individual donor. (C) Same as B, colored according to media. (D) Bray-Curtis distances calculated for each individual and between individuals’ samples and for the same medium or different media. (E) Relative abundances for *B. fragilis, R. gnavus* and *A. muciniphila*.

**Supplementary Figure 2.**
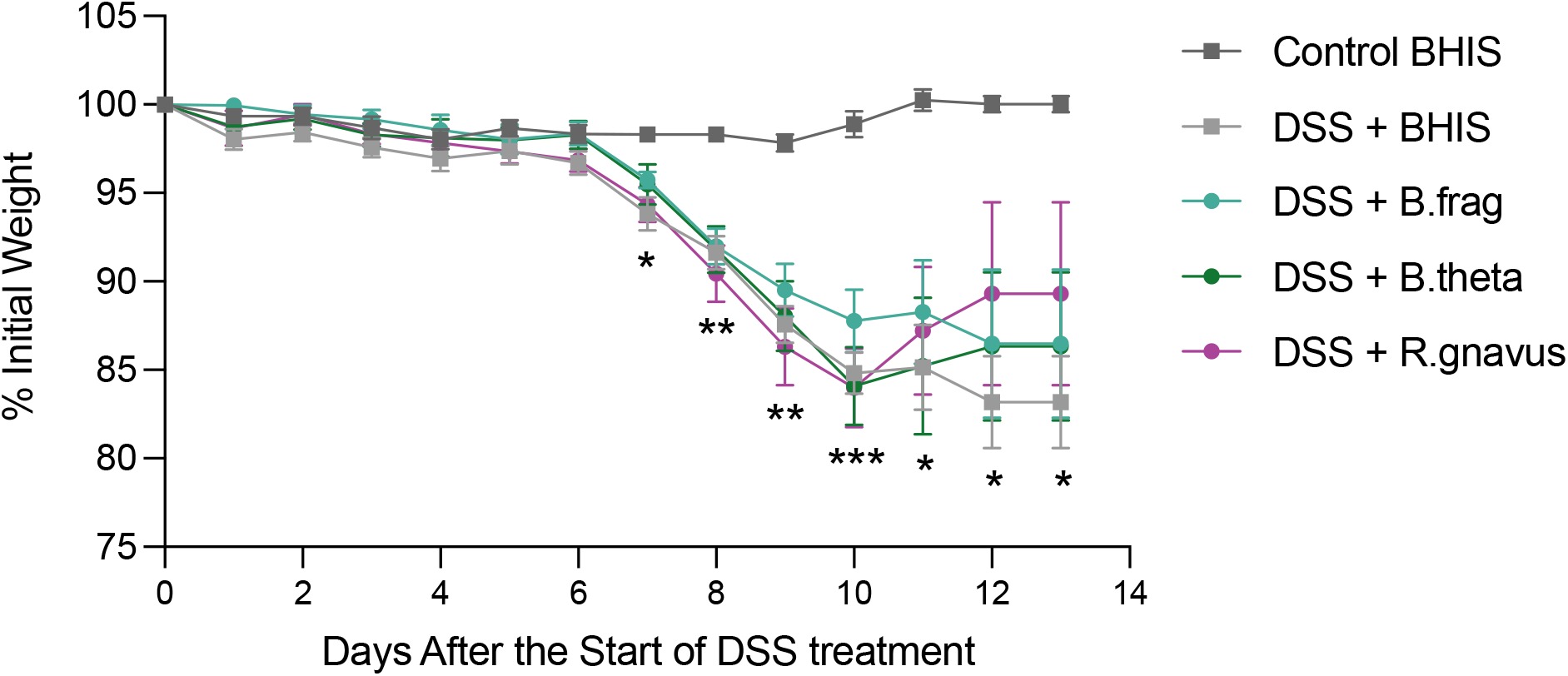
No significant weight loss differences were observed between treatment groups in a DSS-induced mouse model of IBD. Weight loss % after the start of DSS treatment. Data represent mean ± SD. *p<0.05, **p<0.01, and ***p<0.001 for all experimental groups compared to Control BHIS. Statistical significance was assessed using a mixed-effects model with the Geisser-Greenhouse correction followed by Tukey’s multiple comparisons test.

## References

1. Mortensen JH, Lindholm M, Langholm LL, Kjeldsen J, Bay-Jensen AC, Karsdal MA, Manon-Jensen T. 2019. The intestinal tissue homeostasis – the role of extracellular matrix remodeling in inflammatory bowel disease. https://doi.org/101080/1747412420191673729 13:977–993.

2. Mortensen JH, Manon-Jensen T, Jensen MD, Hägglund P, Klinge LG, Kjeldsen J, Krag A, Karsdal MA, Bay-Jensen A-C. 2017. Ulcerative colitis, Crohn’s disease, and irritable bowel syndrome have different profiles of extracellular matrix turnover, which also reflects disease activity in Crohn’s disease. PLoS One 12:e0185855.

3. Petrey AC, de la Motte CA. 2017. The extracellular matrix in IBD. Curr Opin Gastroenterol 33:234– 238.

4. Shimshoni E, Yablecovitch D, Baram L, Dotan I, Sagi I. 2015. ECM remodelling in IBD: Innocent bystander or partner in crime? The emerging role of extracellular molecular events in sustaining intestinal inflammation. Gut 64:367–372.

5. van Haaften WT, Mortensen JH, Karsdal MA, Bay-Jensen AC, Dijkstra G, Olinga P. 2017. Misbalance in type III collagen formation/degradation as a novel serological biomarker for penetrating (Montreal B3) Crohn’s disease. Aliment Pharmacol Ther 46:26–39.

6. Lindholm M, Manon-Jensen T, Madsen GI, Krag A, Karsdal MA, Kjeldsen J, Mortensen JH. 2019. Extracellular Matrix Fragments of the Basement Membrane and the Interstitial Matrix Are Serological Markers of Intestinal Tissue Remodeling and Disease Activity in Dextran Sulfate Sodium Colitis. Dig Dis Sci 64:3134–3142.

7. Koelink PJ, Overbeek SA, Braber S, Morgan ME, Henricks PAJ, Roda MA, Verspaget HW, Wolfkamp SC, Te Velde AA, Jones CW, Jackson PL, Blalock JE, Sparidans RW, Kruijtzer JAW, Garssen J, Folkerts G, Kraneveld AD. 2014. Collagen degradation and neutrophilic infiltration: a vicious circle in inflammatory bowel disease. Gut 63:578–587.

8. Kirov S, Sasson A, Zhang C, Chasalow S, Dongre A, Steen H, Stensballe A, Andersen V, Birkelund S, Bennike TB. 2019. Degradation of the extracellular matrix is part of the pathology of ulcerative colitis. Mol Omi 15:67–76.

9. Kirkegaard T, Hansen A, Bruun E, Brynskov J. 2004. Expression and localisation of matrix metalloproteinases and their natural inhibitors in fistulae of patients with Crohn’s disease. Gut 53:701–709.

10. Baugh MD, Perry MJ, Hollander AP, Davies DR, Cross SS, Lobo AJ, Taylor CJ, Evans GS. 1999. Matrix metalloproteinase levels are elevated in inflammatory bowel disease. Gastroenterology 117:814–822.

11. Gundersen MD, Goll R, Fenton CG, Anderssen E, Sørbye SW, Florholmen JR, Paulssen RH. 2019. Fibrosis Mediators in the Colonic Mucosa of Acute and Healed Ulcerative Colitis. Clin Transl Gastroenterol 10:e00082.

12. Mager R, Roda G, Shalaby MK, Vetrano S. 2020. Fibrotic Strictures in Crohn’s Disease: Mechanisms and Predictive Factors. Curr Drug Targets 22:241–251.

13. Lawrance IC, Rogler G, Bamias G, Breynaert C, Florholmen J, Pellino G, Reif S, Speca S, Latella G. 2015. Cellular and Molecular Mediators of Intestinal Fibrosis. J Crohn’s Colitis 11:j.crohns.2014.09.008.

14. Stenke E, Bourke B, Knaus U. 2017. Crohn’s Strictures—Moving Away from the Knife. Front Pediatr 5:141.

15. Mak JWY, Ng SC. 2020. Epidemiology of fibrostenosing inflammatory bowel disease. J Dig Dis 21:332–335.

16. Schwartz DA, Tagarro I, Carmen Díez M, Sandborn WJ. 2019. Prevalence of Fistulizing Crohn’s Disease in the United States: Estimate From a Systematic Literature Review Attempt and Population-Based Database Analysis. Inflamm Bowel Dis 25:1773–1779.

17. Mortensen JH ø., Godskesen LE lbjer., Jensen MD a., Van Haaften WT obia., Klinge LG abriel., Olinga P, Dijkstra G, Kjeldsen J, Karsdal MA sse., Bay-Jensen AC, Krag A. 2015. Fragments of Citrullinated and MMP-degraded Vimentin and MMP-degraded Type III Collagen Are Novel Serological Biomarkers to Differentiate Crohn’s Disease from Ulcerative Colitis. J Crohn’s Colitis 9:863–872.

18. Annaházi A, Molnár T, Farkas K, Rosztóczy A, Izbéki F, Gecse K, Inczefi O, Nagy F, Földesi I, Sz} M, Dabek M, Ferrier L, Theodorou V, Bueno L, Wittmann T, Róka R. 2013. Fecal MMP-9: A New Noninvasive Differential Diagnostic and Activity Marker in Ulcerative Colitis https://doi.org/10.1002/ibd.22996.

19. Manfredi MA, Zurakowski D, Rufo PA, Walker TR, Fox VL, Moses MA. 2008. Increased incidence of urinary matrix metalloproteinases as predictors of disease in pediatric patients with inflammatory bowel disease. Inflamm Bowel Dis 14:1091–1096.

20. Shimshoni E, Adir I, Afik R, Solomonov I, Shenoy A, Adler M, Puricelli L, Sabino F, Savickas S, Mouhadeb O, Gluck N, Fishman S, Werner L, Salame TM, Shouval DS, Varol C, auf dem Keller U, Podestà A, Geiger T, Milani P, Alon U, Sagi I. 2021. Distinct extracellular–matrix remodeling events precede symptoms of inflammation. Matrix Biol 96:47–68.

21. Galipeau HJ, Caminero A, Turpin W, Bermudez-Brito M, Santiago A, Libertucci J, Constante M, Raygoza Garay JA, Rueda G, Armstrong S, Clarizio A, Smith MI, Surette MG, Bercik P, Croitoru K, Verdu EF, Beck P, Bernstein C, Croitoru K, Dieleman L, Feagan B, Griffiths A, Guttman D, Jacobson K, Kaplan G, Krause DO, Madsen K, Marshall J, Moayyedi P, Ropeleski M, Seidman E, Silverberg M, Snapper S, Stadnyk A, Steinhart H, Surette M, Turner D, Walters T, Vallance B, Aumais G, Bitton A, Cino M, Critch J, Denson L, Deslandres C, El-Matary W, Herfarth H, Higgins P, Huynh H, Hyams J, Mack D, McGrath J, Otley A, Panancionne R. 2020. Novel Fecal Biomarkers That Precede Clinical Diagnosis of Ulcerative Colitis. Gastroenterology 0.

22. Desai MS, Seekatz AM, Koropatkin NM, Kamada N, Hickey CA, Wolter M, Pudlo NA, Kitamoto S, Terrapon N, Muller A, Young VB, Henrissat B, Wilmes P, Stappenbeck TS, N????ez G, Martens EC. 2016. A Dietary Fiber-Deprived Gut Microbiota Degrades the Colonic Mucus Barrier and Enhances Pathogen Susceptibility. Cell 167:1339-1353.e21.

23. Tailford LE, Crost EH, Kavanaugh D, Juge N. 2015. Mucin glycan foraging in the human gut microbiome. Front Genet 6:81.

24. Raimondi S, Musmeci E, Candeliere F, Amaretti A, Rossi M. 2021. Identification of mucin degraders of the human gut microbiota. Sci Reports 2021 111 11:1–10.

25. Kamphuis JBJ, Mercier-Bonin M, Eutamène H, Theodorou V. 2017. Mucus organisation is shaped by colonic content; a new view. Sci Rep 7:8527.

26. Johansson MEV, Holmén Larsson JM, Hansson GC. 2011. The two mucus layers of colon are organized by the MUC2 mucin, whereas the outer layer is a legislator of host–microbial interactions. Proc Natl Acad Sci 108:4659–4665.

27. Van Herreweghen F, De Paepe K, Roume H, Kerckhof FM, Van de Wiele T. 2018. Mucin degradation niche as a driver of microbiome composition and Akkermansia muciniphila abundance in a dynamic gut model is donor independent. FEMS Microbiol Ecol 94:186.

28. Tomlin H, Piccinini AM. 2018. A complex interplay between the extracellular matrix and the innate immune response to microbial pathogens. Immunology 155:186–201.

29. Singh B, Fleury C, Jalalvand F, Riesbeck K. 2012. Human pathogens utilize host extracellular matrix proteins laminin and collagen for adhesion and invasion of the host. FEMS Microbiol Rev 36:1122–1180.

30. Janoir C, Péchiné S, Grosdidier C, Collignon A. 2007. Cwp84, a surface-associated protein of Clostridium difficile, is a cysteine protease with degrading activity on extracellular matrix proteins. J Bacteriol 189:7174–7180.

31. Fletcher JR, Pike CM, Parsons RJ, Rivera AJ, Foley MH, McLaren MR, Montgomery SA, Theriot CM. 2021. Clostridioides difficile exploits toxin-mediated inflammation to alter the host nutritional landscape and exclude competitors from the gut microbiota. Nat Commun 2021 121 12:1–14.

32. Fouillen A, Grenier D, Barbeau J, Baron C, Moffatt P, Nanci A. 2019. Selective bacterial degradation of the extracellular matrix attaching the gingiva to the tooth. Eur J Oral Sci 127:313– 322.

33. Iwasaki M, Usui M, Ariyoshi W, Nakashima K, Nagai-Yoshioka Y, Inoue M, Kobayashi K, Nishihara T. 2021. Evaluation of the ability of the trypsin-like peptidase activity assay to detect severe periodontitis. PLoS One 16:e0256538.

34. Marre AT de O, Domingues RMCP, Lobo LA. 2020. Adhesion of anaerobic periodontal pathogens to extracellular matrix proteins. Brazilian J Microbiol 2020 514 51:1483–1491.

35. Hickey CA, Kuhn KA, Donermeyer DL, Porter NT, Jin C, Cameron EA, Jung H, Kaiko GE, Wegorzewska M, Malvin NP, Glowacki RWP, Hansson GC, Allen PM, Martens EC, Stappenbeck TS. 2015. Colitogenic Bacteroides thetaiotaomicron antigens access host immune cells in a sulfatase-dependent manner via outer membrane vesicles. Cell Host Microbe 17:672–680.

36. Benjdia A, Martens EC, Gordon JI, Berteau O. 2011. Sulfatases and a radical S-adenosyl-L-methionine (AdoMet) enzyme are key for mucosal foraging and fitness of the prominent human gut symbiont, Bacteroides thetaiotaomicron. J Biol Chem 286:25973–25982.

37. Moncrief JS, Obiso R, Barroso LA, Kling JJ, Wright RL, Van Tassell RL, Lyerly DM, Wilkins TD. 1995. The enterotoxin of Bacteroides fragilis is a metalloprotease. Infect Immun 63:175–181.

38. Sánchez E, Laparra JM, Sanz Y. 2012. Discerning the role of bacteroides fragilis in celiac disease pathogenesis. Appl Environ Microbiol 78:6507–6515.

39. Durant L, Stentz R, Noble A, Brooks J, Gicheva N, Reddi D, O’Connor MJ, Hoyles L, McCartney AL, Man R, Pring ET, Dilke S, Hendy P, Segal JP, Lim DNF, Misra R, Hart AL, Arebi N, Carding SR, Knight SC. 2020. Bacteroides thetaiotaomicron-derived outer membrane vesicles promote regulatory dendritic cell responses in health but not in inflammatory bowel disease. Microbiome 8:1–16.

40. Png CW, Lindén SK, Gilshenan KS, Zoetendal EG, McSweeney CS, Sly LI, McGuckin MA, Florin THJ. 2010. Mucolytic Bacteria With Increased Prevalence in IBD Mucosa Augment In Vitro Utilization of Mucin by Other Bacteria. Am J Gastroenterol 105:2420–2428.

41. Danilova NA, Александровна ДН, Abdulkhakov SR, Рустамович АС, Grigoryeva T V, Владимировна ГТ, Markelova MI, Ивановна ММ, Vasilyev IY, Юрьевич В И, Boulygina EA, Александровна БЕ, Ardatskaya MD, Дмитриевна АМ, Pavlenko A V, Владимирович ПА, Tyakht A V, Викторович ТА, Odintsova AK, Харисовна ОА, Abdulkhakov RA, Аббасович АР. 2019. Markers of dysbiosis in patients with ulcerative colitis and Crohn’s disease. Ter Arkh 91:13–20.

42. Hall AB, Yassour M, Sauk J, Garner A, Jiang X, Arthur T, Lagoudas GK, Vatanen T, Fornelos N, Wilson R, Bertha M, Cohen M, Garber J, Khalili H, Gevers D, Ananthakrishnan AN, Kugathasan S, Lander ES, Blainey P, Vlamakis H, Xavier RJ, Huttenhower C. 2017. A novel Ruminococcus gnavus clade enriched in inflammatory bowel disease patients. Genome Med 9:1–12.

43. Holton J. 2008. Enterotoxigenic Bacteroides fragilis. Curr Infect Dis Reports 2008 102 10:99–104.

44. Antalis TM, Shea-Donohue T, Vogel SN, Sears C, Fasano A. 2007. Mechanisms of Disease: protease functions in intestinal mucosal pathobiology. Nat Clin Pract Gastroenterol Hepatol 2007 47 4:393–402.

45. Zhang C, Yu Z, Zhao J, Zhang H, Zhai Q, Chen W. 2019. Colonization and probiotic function of Bifidobacterium longum. J Funct Foods 53:157–165.

46. Jensen H, Grimmer S, Naterstad K, Axelsson L. 2012. In vitro testing of commercial and potential probiotic lactic acid bacteria. Int J Food Microbiol 153:216–222.

47. Brito IL, Yilmaz S, Huang K, Xu L, Jupiter SD, Jenkins AP, Naisilisili W, Tamminen M, Smillie CS, Wortman JR, Birren BW, Xavier RJ, Blainey PC, Singh AK, Gevers D, Alm EJ. 2016. Mobile genes in the human microbiome are structured from global to individual scales. Nature 535:435–9.

48. Goodman AL, Kallstrom G, Faith JJ, Reyes A, Moore A, Dantas G, Gordon JI. 2011. Extensive personal human gut microbiota culture collections characterized and manipulated in gnotobiotic mice. Proc Natl Acad Sci 108:6252–6257.

49. Chassaing B, Srinivasan G, Delgado MA, Young AN, Gewirtz AT, Vijay-Kumar M. 2012. Fecal Lipocalin 2, a Sensitive and Broadly Dynamic Non-Invasive Biomarker for Intestinal Inflammation. PLoS One 7:44328.

50. Franzosa EA, Sirota-Madi A, Avila-Pacheco J, Fornelos N, Haiser HJ, Reinker S, Vatanen T, Hall AB, Mallick H, McIver LJ, Sauk JS, Wilson RG, Stevens BW, Scott JM, Pierce K, Deik AA, Bullock K, Imhann F, Porter JA, Zhernakova A, Fu J, Weersma RK, Wijmenga C, Clish CB, Vlamakis H, Huttenhower C, Xavier RJ. 2018. Gut microbiome structure and metabolic activity in inflammatory bowel disease. Nat Microbiol 2018 42 4:293–305.

51. Derkacz A, Olczyk P, Olczyk K, Komosinska-Vassev K. 2021. The Role of Extracellular Matrix Components in Inflammatory Bowel Diseases. J Clin Med 2021, Vol 10, Page 1122 10:1122.

52. Golusda L, Kühl AA, Siegmund B, Paclik D. 2021. Extracellular Matrix Components as Diagnostic Tools in Inflammatory Bowel Disease. Biol 2021, Vol 10, Page 1024 10:1024.

53. Zhihua L, Aloulou A, Rhimi M, Jablaoui A, Kriaa A, Mkaouar H, Akermi N, Soussou S, Wysocka M, Wołoszyn D, Amouri A, Gargouri A, Maguin E, Lesner A. 2020. Fecal Serine Protease Profiling in Inflammatory Bowel Diseases. Fecal Serine Protease Profiling Inflamm Bowel Dis Front Cell Infect Microbiol 10:21.

54. Mills RH, Dulai PS, Vázquez-Baeza Y, Sauceda C, Daniel N, Gerner RR, Batachari LE, Malfavon M, Zhu Q, Weldon K, Humphrey G, Carrillo-Terrazas M, Goldasich LDR, Bryant MK, Raffatellu M, Quinn RA, Gewirtz AT, Chassaing B, Chu H, Sandborn WJ, Dorrestein PC, Knight R, Gonzalez DJ. 2022. Multi-omics analyses of the ulcerative colitis gut microbiome link Bacteroides vulgatus proteases with disease severity. Nat Microbiol 2022 72 7:262–276.

55. Galipeau HJ, Caminero A, Verdu EF. 2021. Increased Bacterial Proteolytic Activity Detected Before Diagnosis of Ulcerative Colitis. Inflamm Bowel Dis 27:e144–e144.

56. Henke MT, Brown EM, Cassilly CD, Vlamakis H, Xavier RJ, Clardy J. 2021. Capsular polysaccharide correlates with immune response to the human gut microbe Ruminococcus gnavus. Proc Natl Acad Sci U S A 118.

57. Chan JL, Wu S, Geis AL, Chan G V., Gomes TAM, Beck SE, Wu X, Fan H, Tam AJ, Chung L, Ding H, Wang H, Pardoll DM, Housseau F, Sears CL. 2018. Non-toxigenic Bacteroides fragilis (NTBF) administration reduces bacteria-driven chronic colitis and tumor development independent of polysaccharide A. Mucosal Immunol 2018 121 12:164–177.

58. Annaházi A, Gecse K, Dabek M, Ait-Belgnaoui A, Rosztóczy A, Róka R, Molnár T, Theodorou V, Wittmann T, Bueno L, Eutamene H. 2009. Fecal proteases from diarrheic-IBS and ulcerative colitis patients exert opposite effect on visceral sensitivity in mice. PAIN® 144:209–217.

59. Pruteanu M, Hyland NP, Clarke DJ, Kiely B, Shanahan F. 2011. Degradation of the extracellular matrix components by bacterial-derived metalloproteases. Inflamm Bowel Dis 17:1189–1200.

60. Steck N, Mueller K, Schemann M, Haller D. Bacterial proteases in IBD and IBS https://doi.org/10.1136/gutjnl-2011-300775.

61. Fang J, Wang H, Zhou Y, Zhang H, Zhou H, Zhang X. 2021. Slimy partners: the mucus barrier and gut microbiome in ulcerative colitis. Exp Mol Med 2021 535 53:772–787.

62. Michielan A, D’Incà R. 2015. Intestinal Permeability in Inflammatory Bowel Disease: Pathogenesis, Clinical Evaluation, and Therapy of Leaky Gut. Mediators Inflamm 2015:1–10.

63. Steck N, Hoffmann M, Sava IG, Kim SC, Hahne H, Tonkonogy SL, Mair K, Krueger D, Pruteanu M, Shanahan F, Vogelmann R, Schemann M, Kuster B, Sartor RB, Haller D. 2011. Enterococcus faecalis metalloprotease compromises epithelial barrier and contributes to intestinal inflammation. Gastroenterology 141:959–971.

64. Buhr JJ, Blaut & M. 2002. Mucosal and Invading Bacteria in Patients with Inflammatory Bowel Disease Compared with Controls. Scand J Gastroenterol 37:1034–1041.

65. Vrakas S, Mountzouris KC, Michalopoulos G, Karamanolis G, Papatheodoridis G, Tzathas C, Gazouli M. 2017. Intestinal Bacteria Composition and Translocation of Bacteria in Inflammatory Bowel Disease. PLoS One 12:e0170034.

66. Shogan BD, Belogortseva N, Luong PM, Zaborin A, Lax S, Bethel C, Ward M, Muldoon JP, Singer M, An G, Umanskiy K, Konda V, Shakhsheer B, Luo J, Klabbers R, Hancock LE, Gilbert J, Zaborina O, Alverdy JC. 2015. Collagen degradation and MMP9 activation by Enterococcus faecalis contribute to intestinal anastomotic leak. Sci Transl Med 7.

67. Petrey AC, De La Motte CA. 2017. The extracellular matrix in IBD: a dynamic mediator of inflammation. Curr Opin Gastroenterol 33:234.

68. de la Motte CA, Kessler SP. 2015. The Role of Hyaluronan in Innate Defense Responses of the Intestine. Int J Cell Biol 2015:1–5.

69. De La Motte C, Nigro J, Vasanji A, Rho H, Kessler S, Bandyopadhyay S, Danese S, Fiocchi C, Stern R. 2009. Platelet-Derived Hyaluronidase 2 Cleaves Hyaluronan into Fragments that Trigger Monocyte-Mediated Production of Proinflammatory Cytokines. Am J Pathol 174:2254–2264.

70. Mendes RS, Atzingen M V., Domingos RF, Vasconcellos SA, Oliveira R, Vieira ML, Nascimento ALTO. 2012. Plasminogen Binding Proteins and Plasmin Generation on the Surface of Leptospira spp.: The Contribution to the Bacteria-Host Interactions. J Biomed Biotechnol 2012:1–17.

71. Stern M, Stern R. 1992. An ELISA-like assay for hyaluronidase and hyaluronidase inhibitors. Matrix 12:397–403.

72. Andrian E, Grenier D, Rouabhia M. 2004. In vitro models of tissue penetration and destruction by Porphyromonas gingivalis. Infect Immun 72:4689–98.

73. Gommeaux J, Cano C, Garcia S, Gironella M, Pietri S, Culcasi M, Pébusque M-J, Malissen B, Dusetti N, Iovanna J, Carrier A. 2007. Colitis and Colitis-Associated Cancer Are Exacerbated in Mice Deficient for Tumor Protein 53-Induced Nuclear Protein 1. Mol Cell Biol 27:2215–2228.

74. Bonfiglio R, Galli F, Varani M, Scimeca M, Borri F, Fazi S, Cicconi R, Mattei M, Campagna G, Schönberger T, Raymond E, Wunder A, Signore A, Bonanno E. 2021. Extensive histopathological characterization of inflamed bowel in the dextran sulfate sodium mouse model with em-phasis on clinically relevant biomarkers and targets for drug development. Int J Mol Sci 22:1–20.

75. Caporaso JG, Lauber CL, Walters WA, Berg-Lyons D, Lozupone CA, Turnbaugh PJ, Fierer N, Knight R. 2011. Global patterns of 16S rRNA diversity at a depth of millions of sequences per sample. Proc Natl Acad Sci U S A 108:4516–4522.

76. Parada AE, Needham DM, Fuhrman JA. 2016. Every base matters: Assessing small subunit rRNA primers for marine microbiomes with mock communities, time series and global field samples. Environ Microbiol 18:1403–1414.

77. Apprill A, McNally S, Parsons R, Weber L. 2015. Minor revision to V4 region SSU rRNA 806R gene primer greatly increases detection of SAR11 bacterioplankton. Aquat Microb Ecol 75:129– 137.

78. Caporaso JG, Kuczynski J, Stombaugh J, Bittinger K, Bushman FD, Costello EK, Fierer N, Pẽa AG, Goodrich JK, Gordon JI, Huttley GA, Kelley ST, Knights D, Koenig JE, Ley RE, Lozupone CA, McDonald D, Muegge BD, Pirrung M, Reeder J, Sevinsky JR, Turnbaugh PJ, Walters WA, Widmann J, Yatsunenko T, Zaneveld J, Knight R. 2010. QIIME allows analysis of high-throughput community sequencing data. Nat Methods. Nature Publishing Group.

79. Bolyen E, Rideout JR, Dillon MR, Bokulich NA, Abnet CC, Al-Ghalith GA, Alexander H, Alm EJ, Arumugam M, Asnicar F, Bai Y, Bisanz JE, Bittinger K, Brejnrod A, Brislawn CJ, Brown CT, Callahan BJ, Caraballo-Rodríguez AM, Chase J, Cope EK, Da Silva R, Diener C, Dorrestein PC, Douglas GM, Durall DM, Duvallet C, Edwardson CF, Ernst M, Estaki M, Fouquier J, Gauglitz JM, Gibbons SM, Gibson DL, Gonzalez A, Gorlick K, Guo J, Hillmann B, Holmes S, Holste H, Huttenhower C, Huttley GA, Janssen S, Jarmusch AK, Jiang L, Kaehler BD, Kang K Bin, Keefe CR, Keim P, Kelley ST, Knights D, Koester I, Kosciolek T, Kreps J, Langille MGI, Lee J, Ley R, Liu YX, Loftfield E, Lozupone C, Maher M, Marotz C, Martin BD, McDonald D, McIver LJ, Melnik A V., Metcalf JL, Morgan SC, Morton JT, Naimey AT, Navas-Molina JA, Nothias LF, Orchanian SB, Pearson T, Peoples SL, Petras D, Preuss ML, Pruesse E, Rasmussen LB, Rivers A, Robeson MS, Rosenthal P, Segata N, Shaffer M, Shiffer A, Sinha R, Song SJ, Spear JR, Swafford AD, Thompson LR, Torres PJ, Trinh P, Tripathi A, Turnbaugh PJ, Ul-Hasan S, van der Hooft JJJ, Vargas F, Vázquez-Baeza Y, Vogtmann E, von Hippel M, Walters W, Wan Y, Wang M, Warren J, Weber KC, Williamson CHD, Willis AD, Xu ZZ, Zaneveld JR, Zhang Y, Zhu Q, Knight R, Caporaso JG. 2019. Reproducible, interactive, scalable and extensible microbiome data science using QIIME 2. Nat Biotechnol. Nature Publishing Group.

80. Callahan BJ, McMurdie PJ, Rosen MJ, Han AW, Johnson AJA, Holmes SP. 2016. DADA2: High-resolution sample inference from Illumina amplicon data. Nat Methods 13:581–583.

81. Yang Y, Anderson E, Zhang S. 2018. Evaluation of six sample preparation procedures for qualitative and quantitative proteomics analysis of milk fat globule membrane. Electrophoresis 39:2332–2339.

82. Zougman A, Selby PJ, Banks RE. 2014. Suspension trapping (STrap) sample preparation method for bottom-up proteomics analysis. Proteomics 14:1006–1000.

83. Harman RM, He MK, Zhang S, VAN DE WALLE GR. 2018. Plasminogen activator inhibitor-1 and tenascin-C secreted by equine mesenchymal stromal cells stimulate dermal fibroblast migration in vitro and contribute to wound healing in vivo. Cytotherapy 20:1061–1076.

84. Hyatt D, Chen G-L, Locascio PF, Land ML, Larimer FW, Hauser LJ. 2010. Prodigal: prokaryotic gene recognition and translation initiation site identification https://doi.org/10.1186/1471-2105-11-119.

85. Drula E, Garron ML, Dogan S, Lombard V, Henrissat B, Terrapon N. 2022. The carbohydrate-active enzyme database: functions and literature. Nucleic Acids Res 50:D571–D577.

86. Buchfink B, Xie C, Huson DH. 2014. Fast and sensitive protein alignment using DIAMOND. Nat Methods 2014 121 12:59–60.

87. Mistry J, Chuguransky S, Williams L, Qureshi M, Salazar GA, Sonnhammer ELL, Tosatto SCE, Paladin L, Raj S, Richardson LJ, Finn RD, Bateman A. 2021. Pfam: The protein families database in 2021. Nucleic Acids Res 49:D412–D419.

88. Bolger AM, Lohse M, Usadel B. 2014. Trimmomatic: a flexible trimmer for Illumina sequence data. Bioinformatics 30:2114–2120.

